# Numerical simulation of the two-locus Wright-Fisher stochastic differential equation with application to approximating transition probability densities

**DOI:** 10.1101/2020.07.21.213769

**Authors:** Zhangyi He, Mark Beaumont, Feng Yu

**Affiliations:** School of Mathematics, University of Bristol, Bristol BS8 1UG, United Kingdom; School of Biological Sciences, University of Bristol, Bristol BS8 1TQ, United Kingdom

**Keywords:** Natural selection, Two linked loci, Wright-Fisher diffusion, Stochastic Taylor scheme, Importance sampling

## Abstract

Over the past decade there has been an increasing focus on the application of the Wright-Fisher diffusion to the inference of natural selection from genetic time series. A key ingredient for modelling the trajectory of gene frequencies through the Wright-Fisher diffusion is its transition probability density function. Recent advances in DNA sequencing techniques have made it possible to monitor genomes in great detail over time, which presents opportunities for investigating natural selection while accounting for genetic recombination and local linkage. However, most existing methods for computing the transition probability density function of the Wright-Fisher diffusion are only applicable to one-locus problems. To address two-locus problems, in this work we propose a novel numerical scheme for the Wright-Fisher stochastic differential equation of population dynamics under natural selection at two linked loci. Our key innovation is that we reformulate the stochastic differential equation in a closed form that is amenable to simulation, which enables us to avoid boundary issues and reduce computational costs. We also propose an adaptive importance sampling approach based on the proposal introduced by Fearnhead (2008) for computing the transition probability density of the Wright-Fisher diffusion between any two observed states. We show through extensive simulation studies that our approach can achieve comparable performance to the method of Fearnhead (2008) but can avoid manually tuning the parameter *ρ* to deliver superior performance for different observed states.

## 1. Introduction

Throughout the history of population genetics, stochastic models have played a significant role in the study of the population evolving under the influences of various demographic and evolutionary forces within species such as genetic drift and natural selection. One of the most popular stochastic models is the Wright-Fisher model, introduced by Fisher (1922) and Wright (1931), which forms the basis for most theoretical and applied research in population genetics to date, including Kingman’s coalescent (Kingman, 1982), Ewens’ sampling formula (Ewens, 1972), Kimura’s work on fixation probabilities (Kimura, 1955) and statistical techniques for inferring demographic and genetic properties of biological populations (*e.g.*, see Tataru et al., 2017, and references therein for the inference of natural selection with the Wright-Fisher model). However, it is intractable to investigate the gene distribution dynamics in evolving populations through the Wright-Fisher model for large population sizes and evolutionary timescales since a prohibitively large amount of storage and computation is required. The analysis of the Wright-Fisher model today is greatly facilitated by its diffusion approximation, commonly known as the Wright-Fisher diffusion, which can be traced back to Kimura (1964). Crow & Kimura (1970), Ewens (2004) and Durrett (2008) provided an excellent theoretical introduction to the Wright-Fisher model and its diffusion approximation.

In recent years, the Wright-Fisher diffusion has already been successfully applied in the population genetic analysis of time series data of allele frequencies (*e.g.*, Bollback et al., 2008; Gutenkunst et al., 2009; Lukić & Hey, 2012; Malaspinas et al., 2012; Gautier & Vitalis, 2013; Steinrücken et al., 2014; Vitalis et al., 2014; Živković et al., 2015; Ferrer-Admetlla et al., 2016; Schraiber et al., 2016; He et al., 2019), which is mainly due to the fact that it captures the essential features of the underlying evolutionary model and provides a concise framework for characterising the gene distribution dynamics, even in complex evolutionary scenarios. The most important ingredient for the population genetic analysis of genetic time series data through the Wright-Fisher diffusion is its transition probability density function, which can completely determine the gene distribution dynamics in evolving populations. Most existing methods for computing the transition probability density function are based on solving the Kolmogorov equations associated with the Wright-Fisher diffusion, such as finite difference methods (*e.g.*, Boitard & Loisel, 2007; Bollback et al., 2008; Gutenkunst et al., 2009; He et al., 2019), finite volume methods (*e.g.*, Zhao et al., 2013), and spectral methods (*e.g.*, Lukić et al., 2011; Song & Steinrücken, 2012; Steinrücken et al., 2013, 2014, 2015; Živković et al., 2015), most of which, however, are limited to one-locus problems.

With recent advances in DNA preparation and sequencing techniques, the increased availability of genetic time series sampled from multiple loci such as the time serial samples of ancient Eurasians (Mathieson et al., 2015; Allentoft et al., 2015) should provide new opportunities for investigating the gene distribution dynamics underlying such samples, possibly under natural selection, by accounting for the genetic variation at multiple loci jointly. However, according to Kopp et al. (2012), it is infeasible to extend most existing approaches to compute the transition probability density function by solving the Kolmogorov equations associated with the Wright-Fisher diffusion for multi-locus problems since such an approach suffers from dramatically increased computational costs and becomes increasingly inaccurate or even numerically unstable with an increase in the dimension of the problem. Boitard & Loisel (2007) employed a finite difference method to obtain a numerical solution for the probability distribution of haplotype frequencies under the Wright-Fisher model by a diffusion approximation for the two-locus problem of the neutral population. To guarantee a numerically stable computation of the solution, such a method requires an appropriate discretisation for the state space, which may depend strongly on the population genetic quantities such as the recombination rate. However, in population genetic analysis, where we have no prior knowledge of demographic and genetic properties of evolving populations, it is intractable to predict whether the results produced with a particular discretisation are accurate or not. Furthermore, such an approach can only be applied to the states in the interior of the state space, and the approximation of the boundary conditions introduce significant errors.

Fortunately, standard diffusion theory establishes an intimate link between partial differential equation (PDE) and stochastic differential equation (SDE) formulations of the Wright-Fisher diffusion (see, *e.g.*, Karlin & Taylor, 1981), which provides an opportunity for investigating the gene distribution dynamics in evolving populations under the Wright-Fisher diffusion by simulating the SDE instead of solving the PDE (*i.e.*, Kolmogorov backward equation or its adjoint). A number of specialised temporal discretisation methods have been developed for simulating the Wright-Fisher diffusion (*e.g.*, Schurz, 1996; Dangerfield et al., 2012; Schraiber et al., 2013; Neuenkirch & Szpruch, 2014). Recently, Jenkins & Spanó (2017) proposed an approach for exact simulation of the Wright-Fisher diffusion. However, all of these methods are still limited to one-locus problems.

In this work, we introduce a Wright-Fisher SDE of population dynamics subject to natural selection at two linked loci, where the genetic recombination effect and local linkage information are explicitly modelled, and propose an efficient numerical scheme for simulating the Wright-Fisher diffusion. The key innovation of our approach is that we reformulate the Wright-Fisher SDE into a closed form that is amenable to simulation, which enables us to avoid boundary issues and reduce computational costs. Stochastic Taylor schemes are used to obtain an approximate, but often very accurate, solution of the Wright-Fisher SDE, and the associated discretisation error can be reduced by letting the discretisation become more refined. Inspired by Fearnhead (2008), we also develop an adaptive importance sampling method for computing the transition probability density of the Wright-Fisher diffusion between any two observed states with a set of simulated sample paths of the Wright-Fisher diffusion. Unlike the method of Fearnhead (2008), our approach can automatically tune the parameter *ρ* for different observed states and achieve comparable performance. Our method can play an important role in many existing simulation-based inference approaches such as simulated maximum-likelihood methods and Markov chain Monte Carlo methods (see Fuchs, 2013, and reference therein).

The remainder of this work is organised as follows. In Section 2, we provide a brief review of the Wright-Fisher model and its diffusion approximation for the population evolving subject to natural selection at two linked loci. In Section 3, we construct a stochastic Taylor scheme to numerically solve the Wright-Fisher SDE. In Section 4, we develop a numerical approximation of the transition probability density of the Wright-Fisher diffusion through importance sampling and compare the performance of several different importance samplers. Finally, in Section 5, we summarise our main results obtained in the previous sections and discuss possible applications and extensions.

## 2. Wright-Fisher diffusion

In this section, we begin with a short review of the Wright-Fisher model for a population of diploid individuals evolving subject to natural selection at a pair of linked loci. We then develop the diffusion approximation of the Wright-Fisher model in both SDE and PDE formulations.

### 2.1. Wright-Fisher model

Let us consider a diploid population of *N* randomly mating individuals at two linked autosomal loci 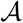 and 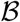 evolving under natural selection with discrete and nonoverlapping generations. We assume that the 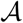 and 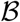 loci are located on a same chromosome with recombination rate *r* ∈ [0, 0.5] between them (*i.e.*, the rate that a recombinant gamete is produced at meiosis), and at each locus there are two allele types, labelled 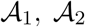 and 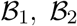, separately, which gives rise to four possible haplotypes 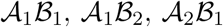 and 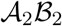, called haplotypes 1, 2, 3 and 4, respectively. We attach the symbols 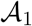 and 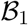 to the mutant alleles, which are assumed to arise only once in the population, and we attach the symbols 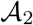 and 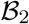 to the ancestral alleles, which are assumed to originally exist in the population.

We incorporate viability selection into the population dynamics and let *w*_*ij*_ represent the viability of an individual with the genotype formed by haplotype *i* and haplotype *j*. In addition to positing absence of sex effects, *i.e.*, *w*_*ij*_ = *w*_*ji*_, we assume that the viability of the genotype at the two loci is determined multiplicatively from the viabilities at individual loci and is fixed from the time that the mutant allele arises. More specifically, the viabilities of the three possible genotypes at each locus, *e.g.*, genotypes 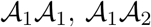 and 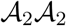 at a given locus 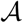, are taken to be 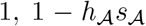 and 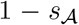, respectively, where 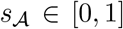 is the selection coefficient and 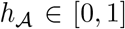 is the dominance parameter. For example, the viability of an individual with the 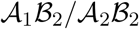 genotype is 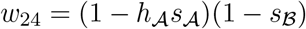.

Let 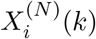 denote the frequency of gamete *i* in a population of *N* individuals in generation 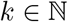 for *i* = 1, 2, 3, 4, and ***X***^(*N*)^(*k*) denote the vector of frequencies of the four possible gametes. In the Wright-Fisher model, the population size *N* is fixed over time, and gametes are chosen randomly from an effectively infinite gamete pool reflecting the parental gamete frequencies at each generation. Under multinomial sampling, we have

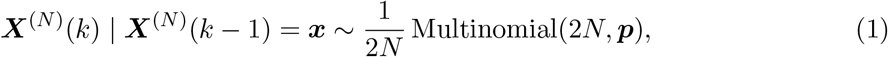

where ***p*** is the vector of the parental gamete frequencies in generation *k* − 1, *i.e.*, the haplotype frequencies of the effectively infinite population after a single generation of random mating, natural selection and genetic recombination. We can express the sampling probabilities ***p*** as a function of the gamete frequencies ***x***,

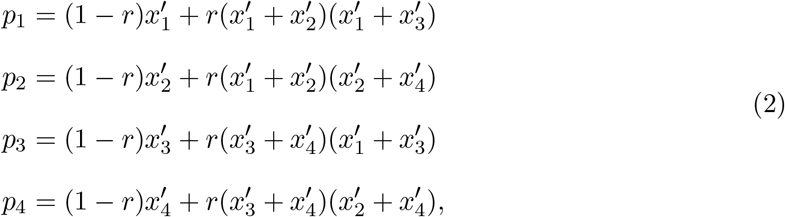

where

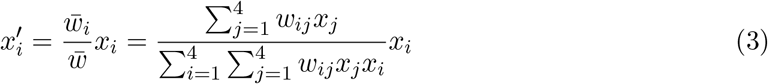

for *i* = 1, 2, 3, 4. In Eq. (3), the numerator 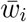 is the marginal fitness of gamete *i* and the denominator 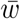 is the mean fitness.

We define the two-locus Wright-Fisher model with selection to be the stochastic process 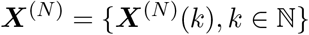 evolving in the state space

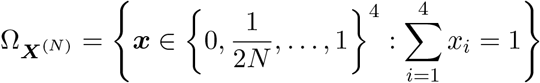

with a multinomial one-step transition probability matrix. From Eqs. (1)-(3), we observe that the transition probabilities of the gamete frequencies from generation *k* to the next are only dependent on the gamete frequencies in generation *k* but are independent of generation *k*, which implies that the Wright-Fisher model is a time-homogeneous Markov process.

### 2.2. Diffusion approximation

Owing to the interplay of stochastic and deterministic forces, the two-locus Wright-Fisher model with selection presents analytical challenges far beyond the Wright-Fisher model for neutral evolution at a single locus. Moreover, it requires a prohibitively large amount of storage and computation to explore the gene distribution dynamics in evolving populations under the Wright-Fisher model when population sizes and evolutionary timescales become large. However, fortunately, we can find an accurate diffusion approximation of the two-locus Wright-Fisher model with selection by running time at rate 2*N*, *i.e.*, *t* = *k*/(2*N*), and holding the scaled selection coefficients 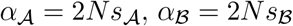 and the scaled recombination rate *ρ* = 4*Nr* fixed while taking the population size *N* → ∞.

#### Theorem 1.

*If we let the population size N go to infinity and hold the scaled parameters* 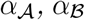 *and R constant, the two-locus Wright-Fisher model with selection **X**^(N)^ converges to a diffusion process, denoted by **X*** = {***X**(t), t* ≥ 0}, *satisfying the stochastic differential equation of the form*

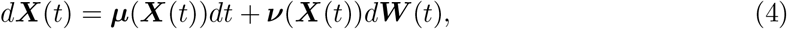

*where the drift term **μ**(**x**) is a four-dimensional vector being*

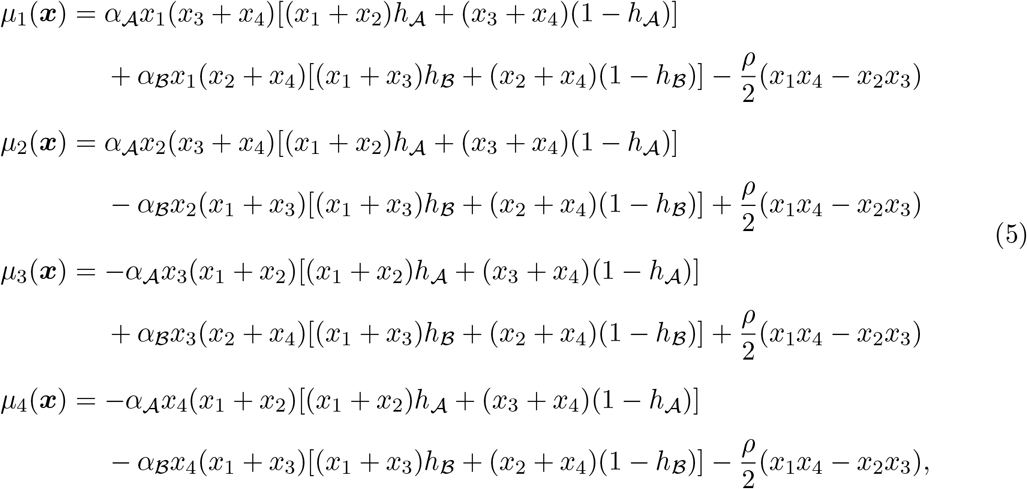

*the diffusion term **ν**(**x**) is a four by three matrix satisfying*

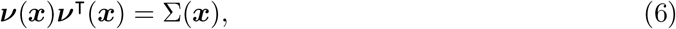

*where*

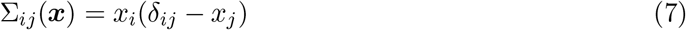

*for i, j* = 1, 2, 3, 4*, and **W*** (*t*) *is a three-dimensional standard Brownian motion.*

The proof of Theorem 1 follows in the similar manner to that employed for neutral populations by Durrett (2008, p. 323), and the major change being the substitution of the gamete frequencies of an effectively infinite population after a single generation of random mating, natural selection and genetic recombination for the sampling probabilities (see He et al., 2017). The term *x*_1_*x*_4_ − *x*_2_*x*_3_ in Eq. (3) is a measure of the linkage disequilibrium between loci 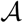 and 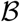, which quantifies the non-random association of the alleles at the two loci. We refer to the diffusion process ***X*** satisfying the SDE in Eq. (4) as the two-locus Wright-Fisher diffusion with selection, which evolves in the state space

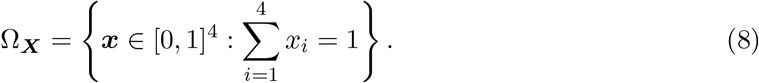

Moreover, for any ***x*** in the interior of the state space Ω_***X***_, the infinitesimal mean vector ***μ***(***x***) and the infinitesimal covariance matrix **Σ**(***x***) are twice continuously differentiable with respect to ***x*** with bounded derivatives, and the infinitesimal covariance matrix **Σ**(***x***) is also positive definite. Therefore, for any ***x*** and ***x′*** in the interior of the state space Ω_***X***_, the transition probability distribution function *P* (*t*, ***x***, ***x′***) of the Wright-Fisher diffusion ***X*** is strongly continuous with respect to Lebesgue’s measure (Stroock & Varadhan, 1979), and the transition probability density function *p*(*t*, ***x***, ***x′***) satisfies the Kolmogorov backward equation (KBE)

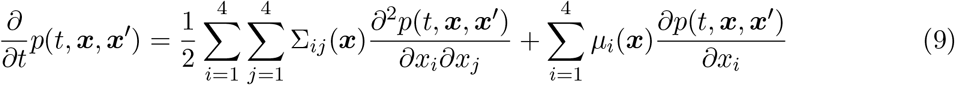

for fixed ***x′*** and the Kolmogorov forward equation (KFE)

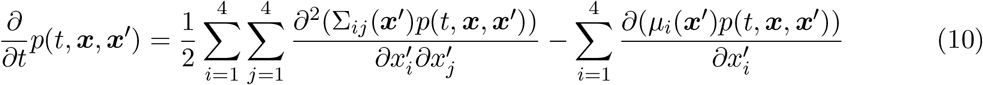

for fixed ***x***, where the infinitesimal mean vector ***μ***(***x***) and the infinitesimal covariance matrix **Σ**(***x***) are given by Eqs. (5) and (7), respectively.

To solve the KBE in Eq. (9) or the KFE in Eq. (10), we need to specify certain boundary conditions. At the boundary of the state space Ω_***X***_, the Wright-Fisher diffusion ***X*** can have different behaviours depending on the type of the boundary: corners, edges and faces. Indeed, its boundary behaviour corresponds to a regular boundary except in the case that fixation occurs at least at one locus (*i.e.*, corners and edges). Since fixation at one locus is non-reversible, this kind of boundary behaviour is characteristic of an exit boundary. When fixation occurs at one locus (*i.e.*, edges), the two-locus Wright-Fisher diffusion with selection reduces to the one-locus Wright-Fisher diffusion with selection, for the population evolving at the other locus. Notice that the classification of boundaries here is extended from Feller’s boundary classification (see Karlin & Taylor, 1981, for more details). Even though the boundary behaviour of the Wright-Fisher diffusion ***X*** is known, it is still intractable to formulate the boundary conditions with exact equations, which could cause a serious problem in solving the KBE and the KFE, either analytically or numerically (see Boitard & Loisel, 2007).

## 3. Stochastic Taylor scheme

Although the Kolmogorov equations satisfied by the transition probability density function of the Wright-Fisher diffusion ***X*** can be readily obtained in Section 2.2, unfortunately, to the best of our knowledge, there are still no known analytical solutions to the Kolmogorov equations, not even numerical solutions. In this section, we resort to a numerical simulation scheme to solve the Wright-Fisher SDE in Eq. (4) rather than the resulting Kolmogorov equations in Eq. (9) or (10).

### 3.1. Equivalent SDE models

To find a numerical solution of the Wright-Fisher SDE in Eq. (4), we need to compute the diffusion term ***σ***(***x***), which we have to perform at every time step in most existing numerical simulation schemes. The diffusion term ***σ***(***x***) can be derived using the Cholesky decomposition. This has been given analytically by Eq. (5.3) in Sato (1976). However, many coefficients in this decomposition explodes at the boundaries. For example,

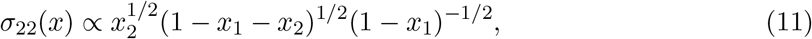

which will be numerically unstable when *x*_1_ is close to 1. More precisely, even though in Eq. (11) (1 − *x*_1_ − *x*_2_)^1/2^ will be tiny, which cancels out the magnitude of (1 − *x*_1_)^−1/2^, any round-off error will be magnified when computing (1 − *x*_1_)^−1/2^. There exist other matrix decompositions capable of computing the diffusion term ***σ***(***x***) such as spectral decomposition, which are valid for positive semi-definite matrices, typically at the expense of either additional numerical errors and computational costs, or limitations in applicability to the infinitesimal covariance matrix **Σ**(***x***) of the form in Eq. (7).

Inspired by Allen et al. (2008), we construct a structurally different SDE from the population evolving subject to natural selection at two linked loci, which is equivalent to the Wright-Fisher SDE in Eq. (4) but enables us to avoid the problem with the decomposition of the infinitesimal covariance matrix **Σ**(***x***). Such an SDE can be formulated in the form:

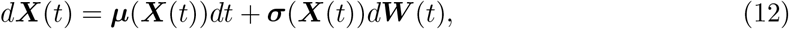

where the diffusion term ***σ***(***x***) can be explicitly written down as

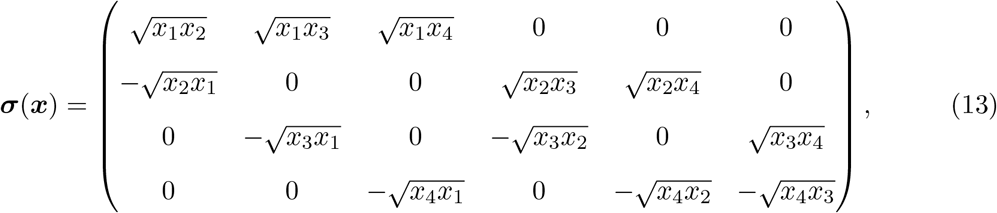

and ***W*** (*t*) is a six-dimensional standard Brownian motion. We observe that unlike the Cholesky decomposition found in Eq. (5.3) of Sato (1976), there is no division or negative power in the diffusion term ***σ***(***x***) in Eq. (13), which can lead to numerical instability.

From Eqs. (5) and (13), we find that the drift term ***μ***(***x***) and the diffusion term ***σ***(***x***) do not satisfy the local Lipschitz conditions (see Karlin & Taylor, 1981), but do satisfy the following conditions arising in the study of the Brownian motion on the group of diffeomorphisms of the circle (*e.g.*, Malliavin, 1999; Airault & Ren, 2002):

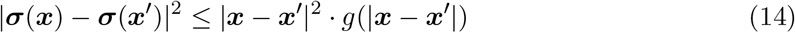

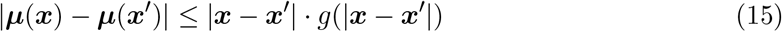

for all ***x***, ***x′*** ∈ Ω_***X***_, where | · | denotes the Euclidean vector norm or the Frobenius matrix norm as appropriate and

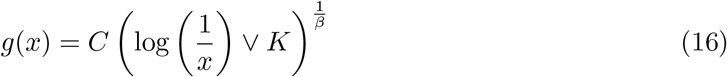

for some *C* > 0, *K* > 0 and *β* > 1. Here we can take *C* = 1, *K* = 1 and *β* = 2, respectively. According to Yamada & Watanabe (1971) and Taniguchi (1992), the non-Lipschitz conditions (14) and (15) are sufficient to ensure the existence and uniqueness of the solution for the SDE in Eq. (12). The equivalence of the Wright-Fisher SDE in Eq. (4) and (12) is given by the following theorem, which guarantees that we can obtain the numerical solution of the SDE in Eq. (4) by solving the SDE in Eq. (12) with numerical simulation schemes.

#### Theorem 2.

*Solutions to the Wright-Fisher SDE in Eq. (4) and (12) possess the same probability law.*

The proof of Theorem 2 can be found in Appendix A.

### 3.2. Euler-Maruyama scheme

We now turn to the development of a numerical simulation scheme for the TLWFDS-SDE with the decomposition of the infinitesimal covariance matrix **Σ**(***x***) we proposed in Section 3.1. Assuming that the closed-form expression of the diffusion coefficient matrix ***σ***(***x***) is available, we consider the construction of stochastic Taylor schemes here. Without loss of generality, we take the initial time point to be 0 and focus our attention on the TLWFDS* model for time *t* on a interval [0, *T*] in the following due to the fact that the Wright-Fisher diffusion ***X*** is time-homogeneous.

To obtain the stochastic Itô-Taylor expansion for the solution of the SDE in Eqs. (11)-(13), we first represent the Wright-Fisher SDE in the integral component form as

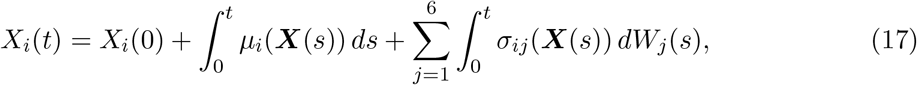

for *i* = 1, 2, 3, 4. Using Itô’s formula, we then have

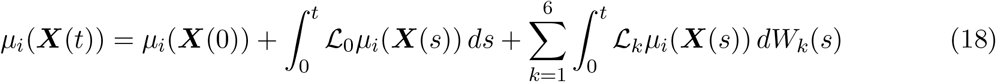

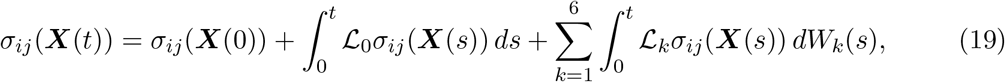

where 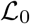 and 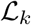 are differential operators defined as

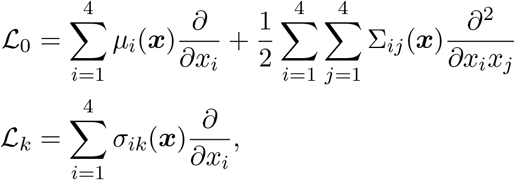

for *k* = 1, 2, …, 6. Substituting Eqs. (18) and (19) into Eq. (17) and separating the constant terms out from the integrand, we can formulate the stochastic Itô-Taylor expansion for the solution of the Wright-Fisher SDE as

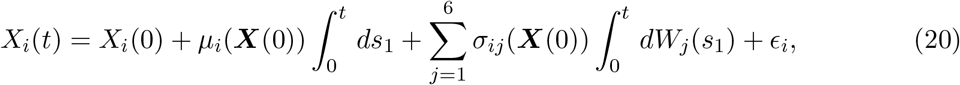

for *i* = 1, 2, 3, 4, where

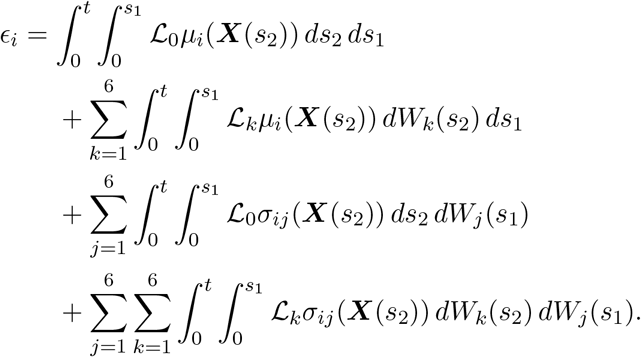

We can get a truncated stochastic Itô-Taylor expansion by keeping the first three constant terms in Eq. (20) or continue applying Itô’s formula to other terms in *ϵ*_*i*_ to include further higher-order constant terms, which enables us to get truncated stochastic Itô-Taylor expansions of higher order but requires more derivatives and integrals. That is, in theory, we can construct the stochastic Taylor scheme of arbitrarily high order by including further components of the stochastic Itô-Taylor expansion for the solution of the Wright-Fisher SDE in Eqs. (11)-(13).

We introduce a partition of the time interval [0, *T*], defined as

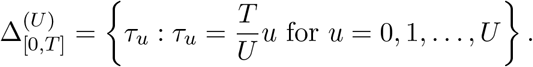

Keeping the first three constant terms in Eq. (20), we can obtain the stochastic Taylor scheme of the form

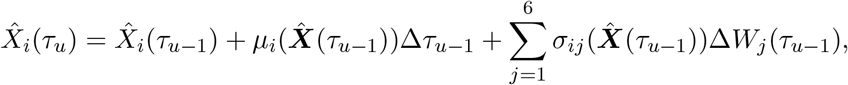

for *i* = 1, 2, 3, 4, where Δ*τ*_*u*−1_ = *τ*_*u*_ − *τ*_*u*−1_ and Δ*W*_*j*_(*τ*_*u*−1_) = *W*_*j*_(*τ*_*u*_)− *W*_*j*_(*τ*_*u*−1_) are independent and normally distributed with mean 0 and variance Δ*τ*_*u*−1_ for *j* = 1, 2, … , 6.

The stochastic Taylor scheme we adopt here is rather simple, commonly known as the Euler-Maruyama scheme, which is one of the most popular stochastic Taylor schemes in practice due to its high efficiency and low complexity. As demonstrated in Kloeden & Platen (1992), the Euler-Maruyama scheme is numerically stable and strongly consistent, and the convergence of the Euler-Maruyama approximation 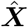 of the Wright-Fisher diffusion is guaranteed by the non-Lipschitz conditions (14) and (15) (see Zhang, 2006, for more details). If one desires a stochastic Taylor scheme of higher order, an efficient systematic procedure proposed by Tocino & Vigo-Aguiar (2003) can be applied to obtain truncated stochastic Itô-Taylor expansions of higher order for the Wright-Fisher diffusion ***X*** expressed in terms of powers of the increments Δ*t* and Δ***X***(*t*).

We illustrate the empirical cumulative distribution functions of the Wright-Fisher diffusion ***X*** for each gamete frequency at the given time point in Figures 1 and 2, which are obtained with a sample of the realizations of the Wright-Fisher diffusion simulated with the Euler-Maruyama scheme. By comparing with the empirical cumulative distribution functions of the Wright-Fisher model ***X***^(*N*)^ for each gamete frequency at the same time point, we find that the Euler-Maruyama approximation 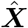 of the Wright-Fisher diffusion is sufficiently accurate on the time interval [0, *T*] for a sufficiently fine partition 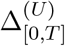 with enough realizations in the sample.

**Figure 1:**
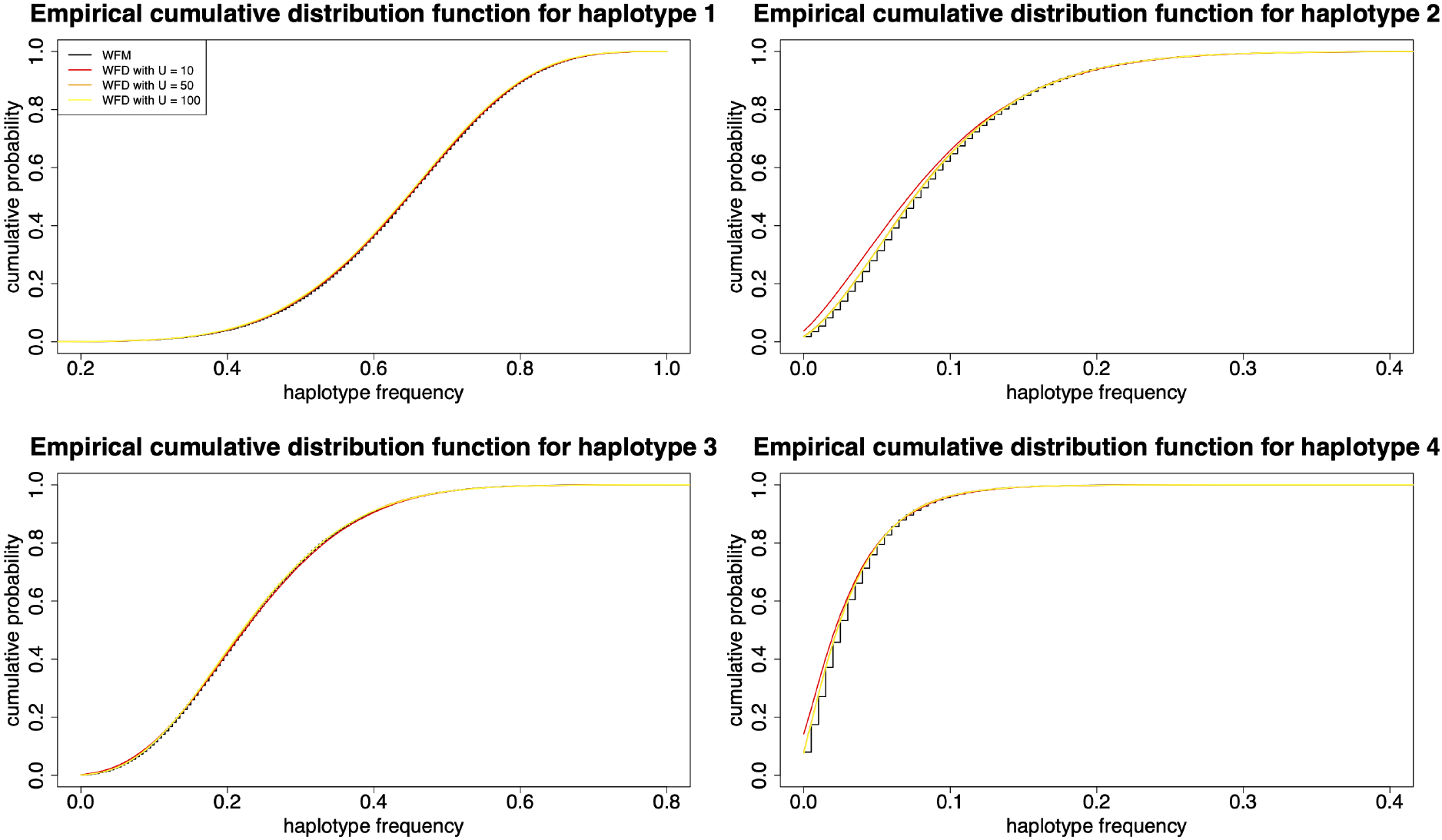
Empirical cumulative distribution functions for each gamete frequency in generation 200 under the Wright-Fisher diffusion ***X*** for different interval partitions. We partition the time interval into *U* = 10, 50, 100 equidistant subintervals for the Euler-Maruyama scheme, respectively, and we construct the empirical cumulative distribution functions with *V* = 100000 independent realizations of the Wright-Fisher diffusion. In addition, we provide the empirical cumulative distribution functions for each haplotype frequency under the Wright-Fisher model ***X***^(*N*)^ for reference. We adopt ***x***_0_ = (0.3, 0.2, 0.2, 0.3), *N* = 1000, 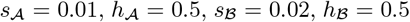 and *r* = 0.05.

**Figure 2:**
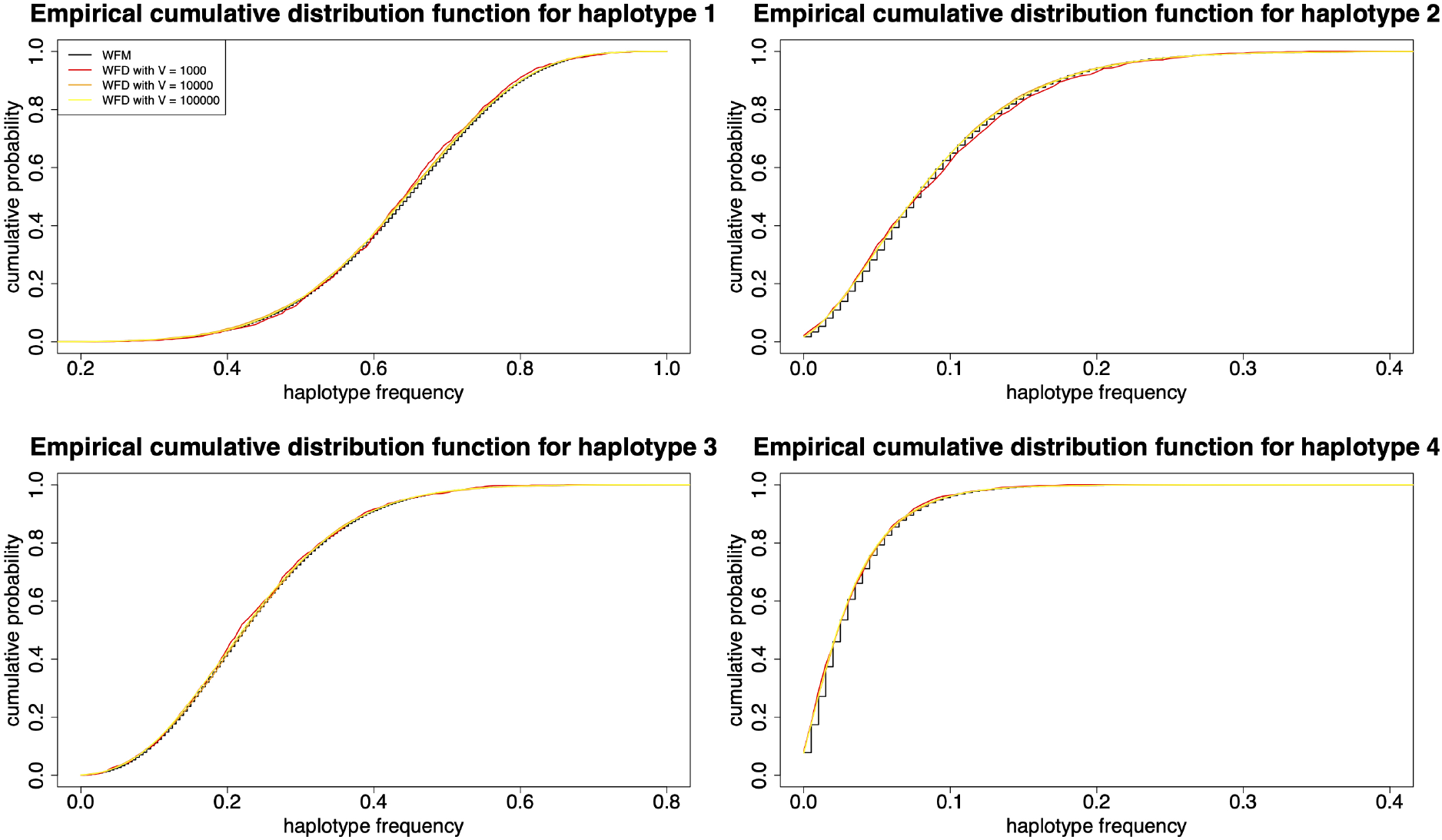
Empirical cumulative distribution functions for each gamete frequency in generation 200 under the Wright-Fisher diffusion ***X*** for different sample sizes. We partition the time interval into *U* = 100 equidistant subintervals for the Euler-Maruyama scheme, and we construct the empirical cumulative distribution functions with *V* = 1000, 10000, 100000 independent realizations of the Wright-Fisher diffusion, respectively. In addition, we provide the empirical cumulative distribution functions for each haplotype frequency under the Wright-Fisher model ***X***^(*N*)^ for reference. We adopt ***x***_0_ = (0.3, 0.2, 0.2, 0.3), *N* = 1000, 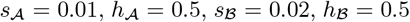 and *r* = 0.05.

## 4. Transition probability densities

As we have stated in Section 1, the most important ingredient in parametric inference procedures for the time series data of allele frequencies under the Wright-Fisher diffusion is the transition probability density function, which can completely characterise the gene distribution dynamics in evolving populations. Most existing approaches for computing the transition probability density function are limited to one-locus problems, and it becomes intractable to extend them to multi-locus problems due to dramatically increased computational costs associated with an increase in the dimensionality of the problem. However, not all existing parametric inference procedures require the transition probability density function (*i.e.*, transition probability densities for all possible states in the state space). For instance, simulated maximum-likelihood methods (*e.g.*, Pedersen, 1995a,b; Brandt & Santa-Clara, 2002; Durham & Gallant, 2002) just need to evaluate the transition probability density of the Wright-Fisher diffusion between two observed states. In this section, we resort to importance sampling techniques to construct an efficient numerical approximation of the transition probability density of the Wright-Fisher diffusion between two observed states, which may introduce some statistical errors but can be reduced by higher-order numerical simulation schemes and larger numbers of simulated sample paths. For simplicity, we demonstrate the construction of the importance sampling approximation of the transition probability density with the Euler-Maruyama scheme in the following.

### 4.1. Euler-Maruyama approximation

Given that the Wright-Fisher diffusion ***X*** is time-homogeneous, we let *p*(*T*, ***x***_0_, ***x***_*T*_) denote the probability density of the transition of the Wright-Fisher diffusion ***X*** from state ***x***_0_ to state ***x***_*T*_, where ***x***_0_ and ***x***_*T*_ are two observed states at the time points 0 and *T*, respectively. If the length of the time interval *T* is sufficiently small, the Euler-Maruyama approximation of the Wright-Fisher diffusion 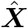 has the desired accuracy on the time interval [0, *T*]. Therefore, we can accurately approximate the transition probability density *p*(*T*, ***x***_0_, ***x***_*T*_) through the Euler-Maruyama approximation of the Wright-Fisher diffusion 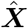 as

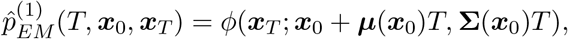

where *ϕ*(·; ***x***_0_ + ***μ***(***x***_0_)*T,* **Σ**(***x***_0_)*T*) is the probability density function of the multivariate normal distribution with the mean vector ***x***_0_ + ***μ***(***x***_0_)*T* and the covariance matrix **Σ**(***x***_0_)*T* . Otherwise, we can use the partition 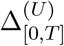 such that the Euler-Maruyama approximation of the Wright-Fisher diffusion 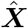 is sufficiently accurate on each subinterval [*τ*_*u*−1_, *τ*_*u*_]. Combining the Markov property of the Wright-Fisher diffusion and the Chapman-Kolmogorov equation, we can accurately approximate the transition probability density *p*(*T*, ***x***_0_, ***x***_*T*_) through the Euler-Maruyama approximation of the Wright-Fisher diffusion 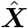 as

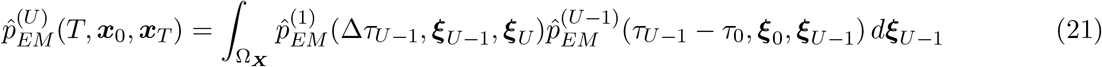

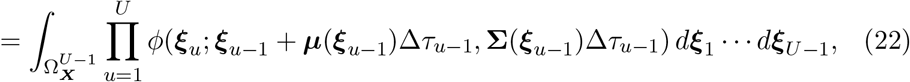

where we define 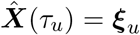 for *u* = 0, 1, … , *U*, and use the convention ***ξ***_0_ = ***x***_0_ and ***ξ***_*U*_ = ***x***_*T*_ to conserve notation.

From Theorem 2 in Pedersen (1995b), we have

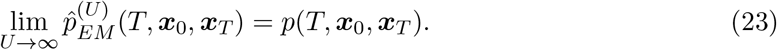

However, it is infeasible to evaluate the integrals in Eq. (22) analytically, and unfortunately, quadrature-based numerical integration techniques become computationally infeasible with a very fine partition of the time interval [0, *T*].

### 4.2. Monte Carlo integration

Inspired by Brandt & Santa-Clara (2002), Monte Carlo integration is an alternative approach of evaluating the integrals in the computation of the Euler-Maruyama approximation 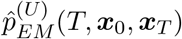. From Eq. (21), using Monte Carlo integration, we can approximate the Euler-Maruyama approximation 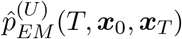 by

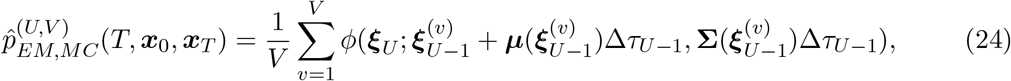

where 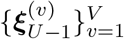 is the sample of the realisations generated by setting ***ξ***_0_ = ***x***_0_ and ***ξ***_*U*_ = ***x***_*T*_, and successively drawing

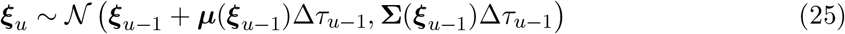

for *u* = 1, 2, … , *U* − 1, and *V* is the total number of the realisations in the sample. In Figure 3, we illustrate a set of the simulated sample paths for the Wright-Fisher diffusion ***X*** with two observed states ***x***_0_ and ***x***_*T*_ generated by using Eq. (25) as an example. However, the simulated sample paths in Figure 3 do not look likely to start at state ***x***_0_ and end at state ***x***_*T*_ .

**Figure 3:**
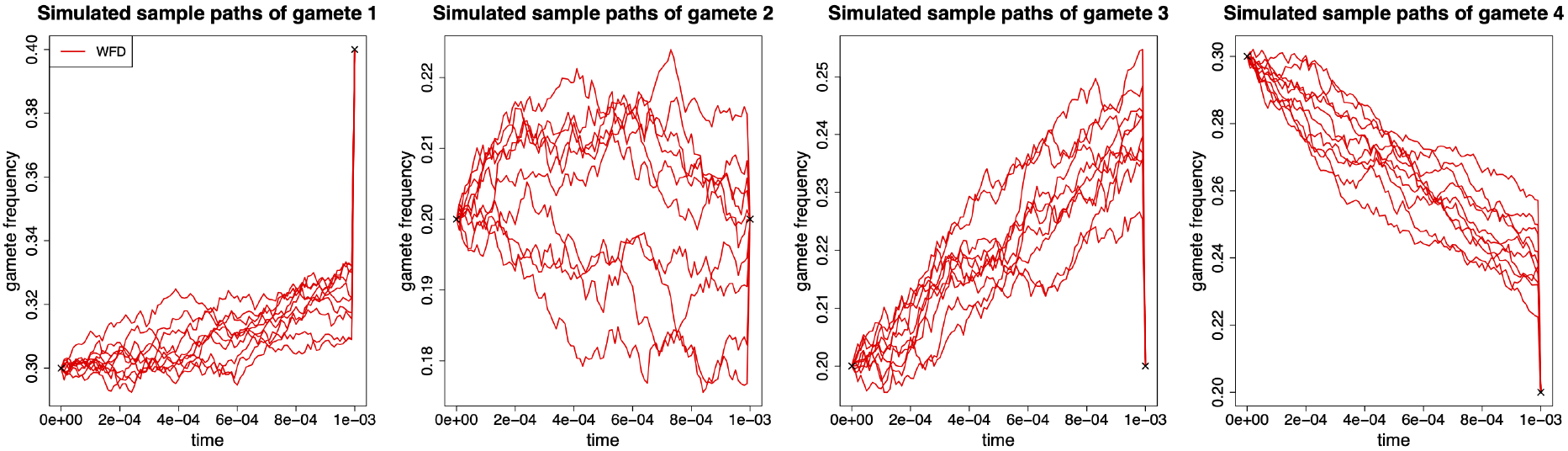
Simulated sample paths of each gamete evolving from generation 0 to 10 drawn from the the Wright-Fisher diffusion. In this illustration, we partition the time interval [0, 0.001] into 100 equidistant subintervals for the Euler-Maruyama scheme and generate 10 simulated sample paths with two observed states ***x***_0_ = (0.3, 0.2, 0.2, 0.3) and ***x***_0.001_ = (0.4, 0.2, 0.2, 0.2). Other parameters *N* = 5000, 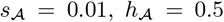, 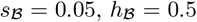 and *r* = 0.05.

The strong law of large numbers guarantees the convergence of the Monte Carlo approximation 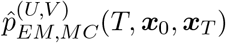 to the Euler-Maruyama approximation 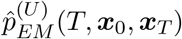 as the sample size *V* goes to infinity. Moreover, combining with Eq. (23), we have

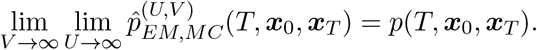

We show the convergence of the Monte Carlo approximation 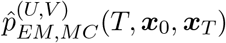 to the transition probability density *p*(*T*, ***x***_0_, ***x***_*T*_) on a logarithmic scale in Figure 4 and find that the Monte Carlo approximation 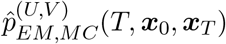 given by Eq. (24) suffers from poor convergence largely due to the enormous variance resulting from Monte Carlo integration. As shown in Figure 3, most of the realisations 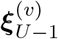 drawn from the Wright-Fisher diffusion ***X*** with the Euler-Maruyama scheme are in regions where the integrand 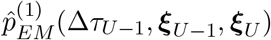 is small, which leads to prohibitive Monte Carlo variability (Durham & Gallant, 2002).

**Figure 4:**
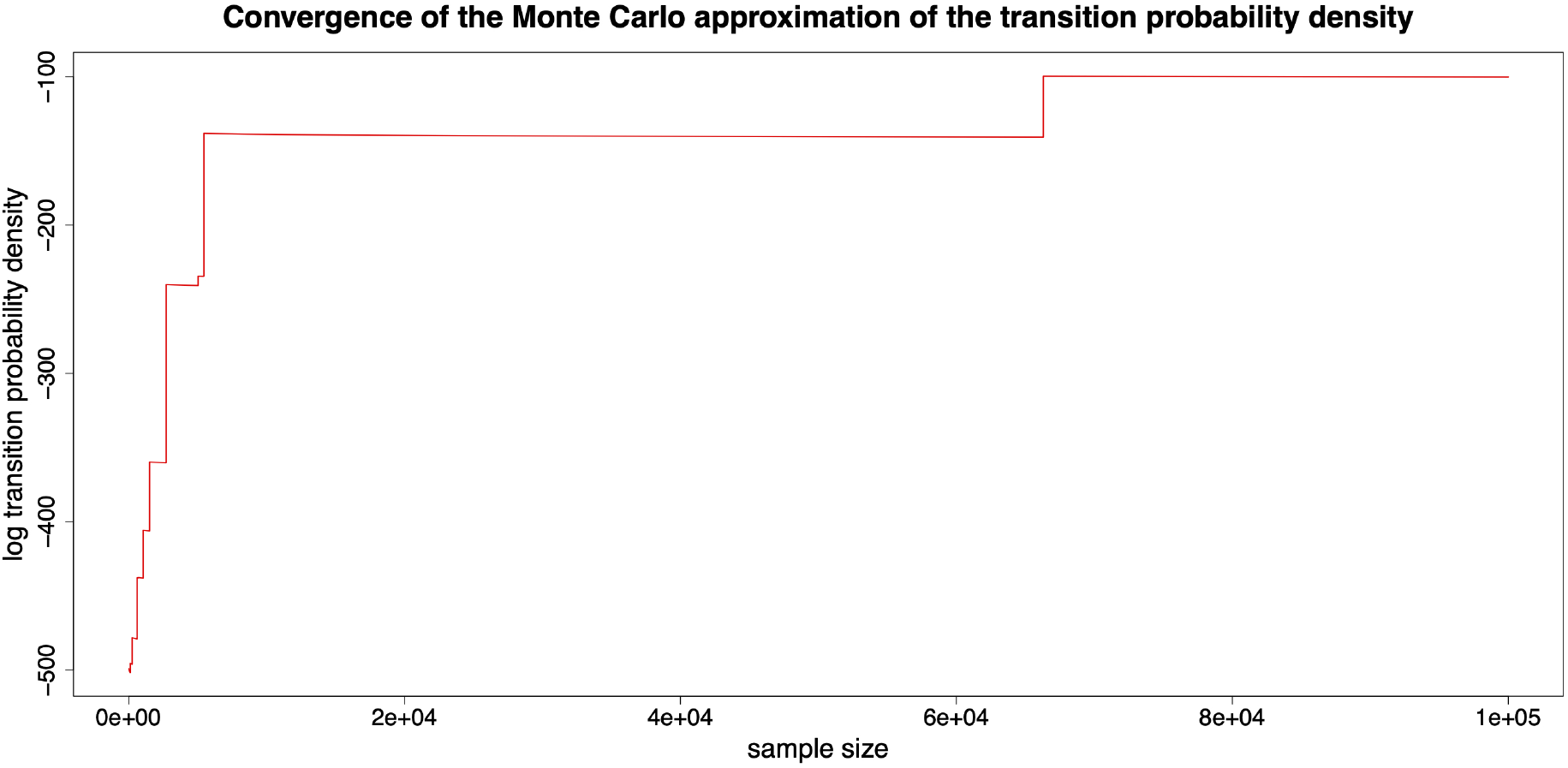
Convergence of the Monte Carlo approximation of the transition probability density 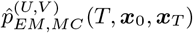. In this illustration, we partition the time interval [0, 0.001] into 100 equidistant subintervals for the Euler-Maruyama scheme and take ***x***_0_ = (0.3, 0.2, 0.2, 0.3), ***x***_0.001_ = (0.4, 0.2, 0.2, 0.2), *N* = 5000, 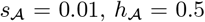, 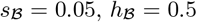 and *r* = 0.05.

### 4.3. Monte Carlo integration with importance sampling

Now we pursue a more efficient Monte Carlo approximation of the transition probability density 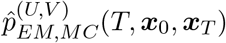 through importance sampling, whose efficiency depends largely on the choice of the importance sampler. Fuchs (2013) provided a number of importance samplers for Monte Carlo variance reduction. In this section, we consider the importance samplers based on the guided process according to the terminology suggested in Papaspiliopoulos & Roberts (2012), which can force the simulated sample paths generated from the importance sampler to terminate at state ***x***_*T*_ almost surely through the so-called guiding drift term defined by

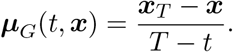

Note that we use *G* to refer to a generic guided process in the following.

The first *G*-based importance sampler we consider was developed by Delyon & Hu (2006), which suggests generating the simulated sample paths from the guided process, denoted by *G*_*1*_, satisfying the following proposal SDE of the form

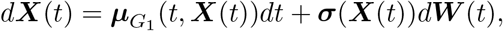

where the drift term

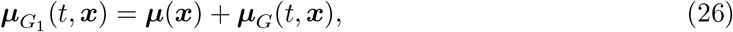

the diffusion term is given by Eq. (13), and ***W*** (*t*) is a six-dimensional standard Brownian motion. From Eq. (26), the drift term 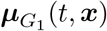 of the *G*_*1*_ is composed of the drift term ***μ***(***x***) of the underlying Wright-Fisher diffusion and the guiding drift term ***μ***_*G*_(*t*, ***x***), which enables the resulting evolution of the *G*_*1*_-SDE to be influenced by both the underlying Wright-Fisher diffusion and the observed state ***x***_*T*_ .

More specifically, using the Euler-Maruyama scheme, we set ***ξ***_0_ = ***x***_0_ and ***ξ***_*U*_ = ***x***_*T*_, and successively draw

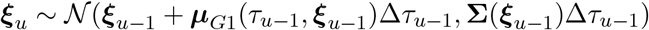

for *u* = 1, 2, … , *U* − 1. We show a set of simulated sample paths generated from the *G*_*1*_-based importance sampler with two observed states ***x***_0_ and ***x***_*T*_ as an example in Figure 5.

**Figure 5:**
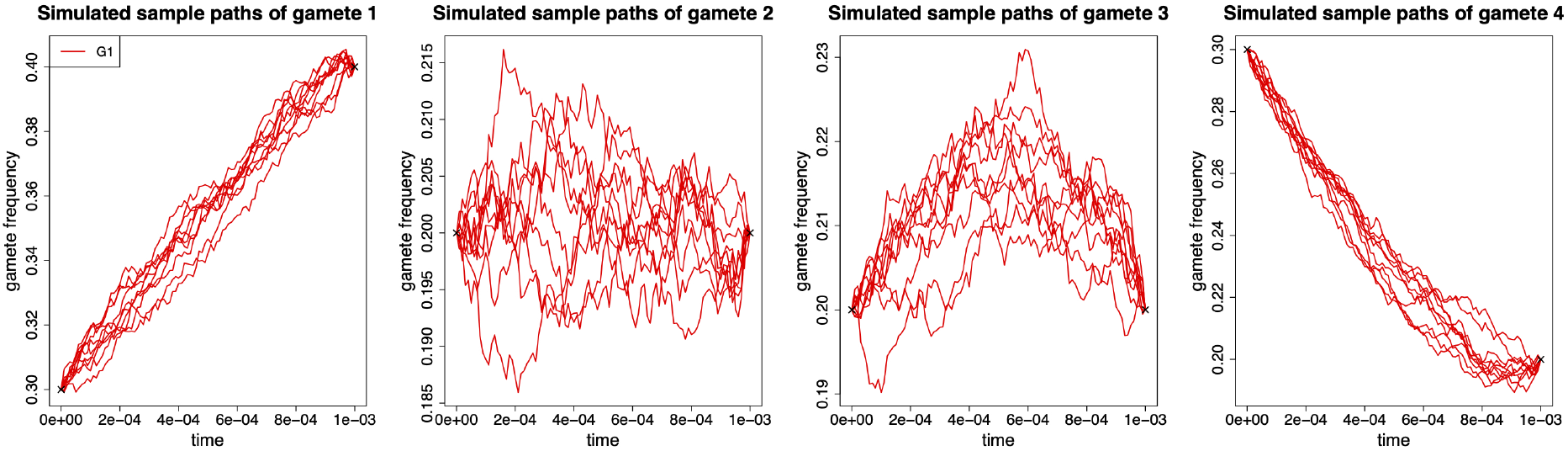
Simulated sample paths of each gamete evolving from generation 0 to 10 drawn from the *G*_*1*_-based importance sampler. In this illustration, we partition the time interval [0, 0.001] into 100 equidistant subintervals for the Euler-Maruyama scheme and generate 10 simulated sample paths with two observed states ***x***_0_ = (0.3, 0.2, 0.2, 0.3) and ***x***_0.001_ = (0.4, 0.2, 0.2, 0.2). Other parameters *N* = 5000, 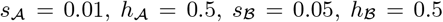 and *r* = 0.05.

**Figure 6:**
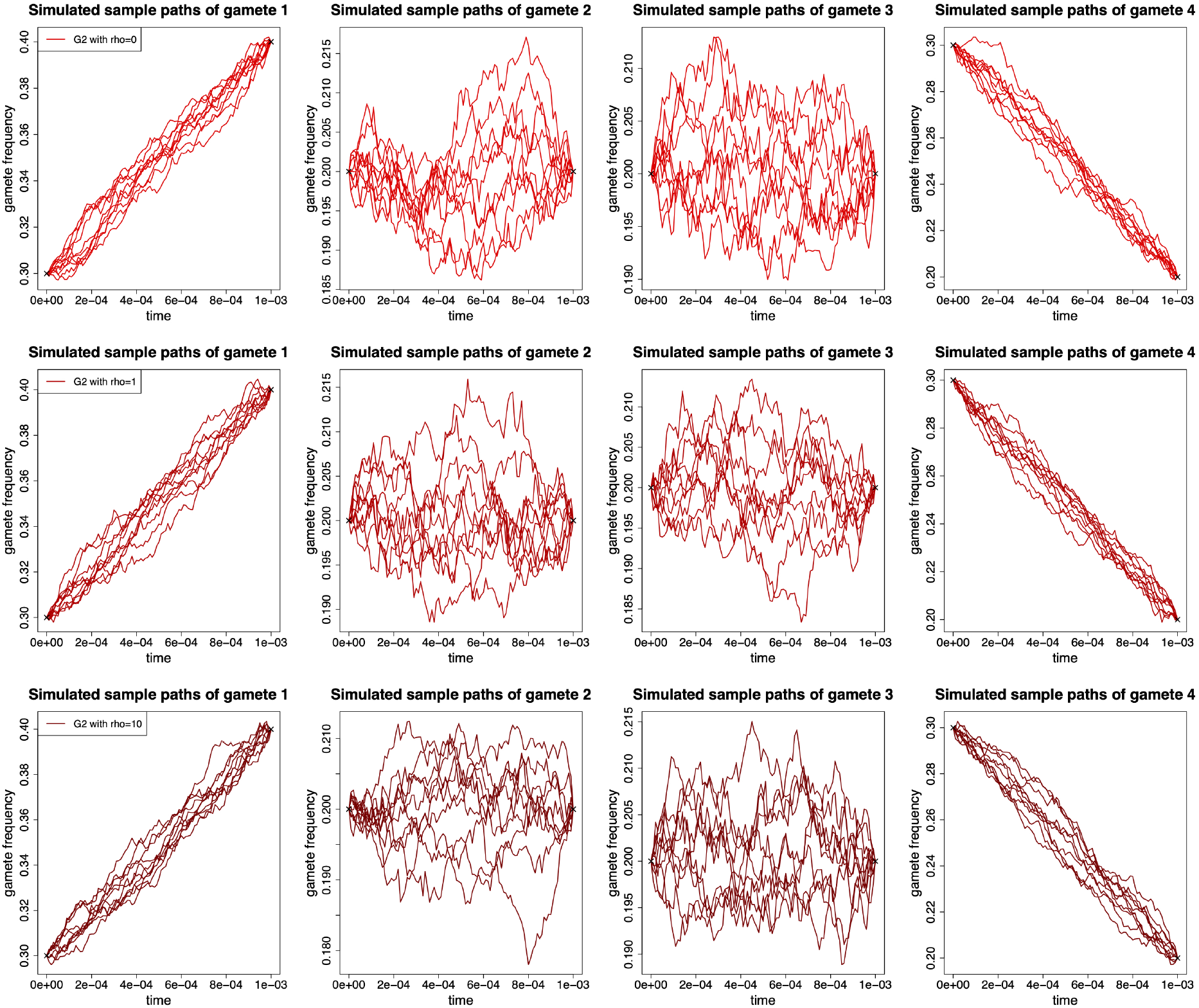
Simulated sample paths of each gamete evolving from generation 0 to 10 drawn from the *G*_*2*_-based importance sampler. In this illustration, we partition the time interval [0, 0.001] into 100 equidistant subintervals for the Euler-Maruyama scheme and generate 10 simulated sample paths with two observed states ***x***_0_ = (0.3, 0.2, 0.2, 0.3) and ***x***_0.001_ = (0.4, 0.2, 0.2, 0.2). Other parameters *N* = 5000, 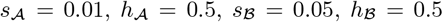 and *r* = 0.05.

The second *G*-based importance sampler we consider was proposed by Fearnhead (2008), which suggests generating the simulated sample paths from the guided process, denoted by *G*_*2*_, satisfying the following proposal SDE of the form

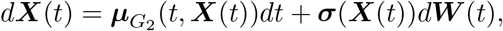

where the drift term

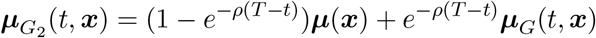

for a predetermined non-negative constant *ρ*, the diffusion term is given by Eq. (13), and ***W*** (*t*) is a six-dimensional standard Brownian motion. Compared to the drift term 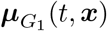 of the *G*_*1*_, the construction of the drift term 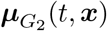 of the *G*_*2*_ increases the contribution of the drift term ***μ***(***x***) of the underlying Wright-Fisher diffusion for time *t* close to 0, whereas for time *t* close to *T*, it increases the contribution of the guiding drift term ***μ***_*G*_(*t*, ***x***).

More specifically, using the Euler-Maruyama scheme, we set ***ξ***_0_ = ***x***_0_ and ***ξ***_*U*_ = ***x***_*T*_, and successively draw

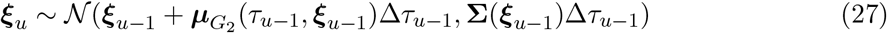

for *u* = 1, 2, … , *U* − 1. In particular, when we set *ρ* = 0 in Eq. (27), the *G*_*2*_-based importance sampler is commonly known as the diffusion bridge proposal. In Figure 7, we illustrate several sets of simulated sample paths generated from the *G*_*2*_-based importance sampler with two observed states ***x***_0_ and ***x***_*T*_ for different values of the constant *ρ* as examples.

**Figure 7:**
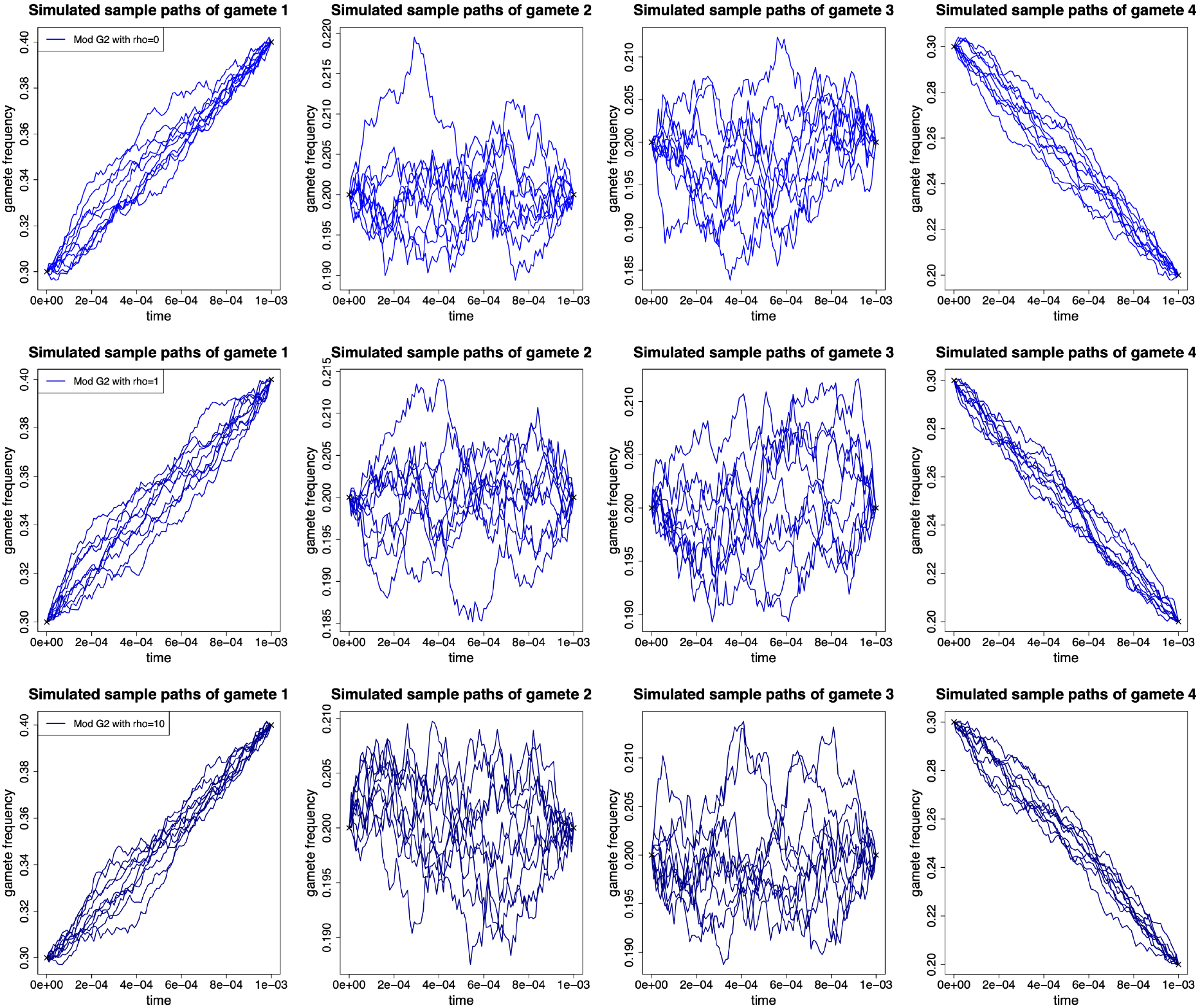
Simulated sample paths of each gamete evolving from generation 0 to 10 drawn from the modified *G*_*2*_-based importance sampler. In this illustration, we partition the time interval [0, 0.001] into 100 equidistant subintervals for the Euler-Maruyama scheme and generate 10 simulated sample paths with two observed states ***x***_0_ = (0.3, 0.2, 0.2, 0.3) and ***x***_0.001_ = (0.4, 0.2, 0.2, 0.2). Other parameters *N* = 5000, 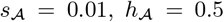, 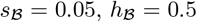 and *r* = 0.05.

Fearnhead (2008) also provided a modified *G*_*2*_-based importance sampler, which suggests successively drawing

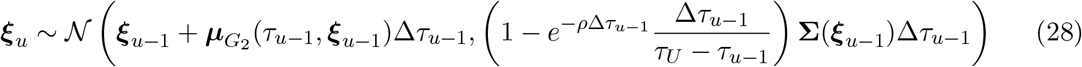

for *u* = 1, 2, … , *U* − 1. When we take *ρ* = 0 in Eq. (28), the modified *G*_*2*_-based importance sampler reduces to the modified diffusion bridge proposal discussed in Durham & Gallant (2002), in which they empirically demonstrate that compared with the diffusion bridge proposal, the modification leads to much better performance in their applications. We illustrate several sets of simulated sample paths generated from the modified *G*_*2*_-based importance sampler with two observed states ***x***_0_ and ***x***_*T*_ for different values of the constant *ρ* as examples in Figure 7.

We propose an adaptive *G*_*2*_-based importance sampler, which suggests generating the simulated sample paths from the guided process, denoted by *G*_*3*_, satisfying the following proposal SDE of the form

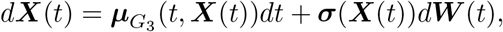

where the drift term

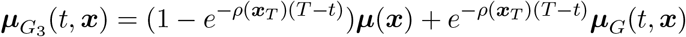

for a predetermined non-negative function *ρ*(***x***_*T*_), the diffusion term is given by Eq. (13), and ***W*** (*t*) is a six-dimensional standard Brownian motion. Compared to the drift term 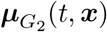 of the *G*_*2*_, the *ρ*-term is a function of the observed state ***x***_*T*_ in the drift term 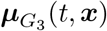 of the *G*_*3*_, which is defined as

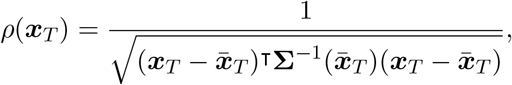

where 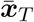 is the value at time *T* of the solution to the following ordinary differential equation (ODE) of the form

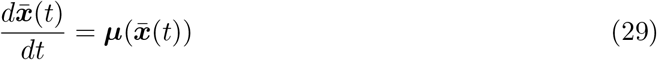

with the initial condition 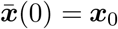. The solution to the ODE (29) characterises an effectively infinite population evolving under natural selection at two linked loci, and therefore 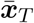 can be considered to be the effectively infinite population limit of ***X***(*T*), where the influence of genetic drift on the changes in gamete frequencies is completely removed. The advantage of constructing such a *ρ*-term is to enable the resulting *G*_*3*_-SDE to evolve largely under the influence of the underlying Wright-Fisher diffusion if its sample path is already close to the observed state ***x***_*T*_ with high probability, but allows a more significant contribution from the guiding drift term if its sample path gets further away from the observed state ***x***_*T*_ .

More specifically, using the Euler-Maruyama scheme, we set ***ξ***_0_ = ***x***_0_ and ***ξ***_*U*_ = ***x***_*T*_, and successively draw

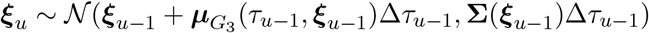

for *u* = 1, 2, … , *U* − 1. Figure 8 illustrates a set of simulated sample paths generated from the *G*_*3*_-based importance sampler with two observed states ***x***_0_ and ***x***_*T*_ as an example.

**Figure 8:**
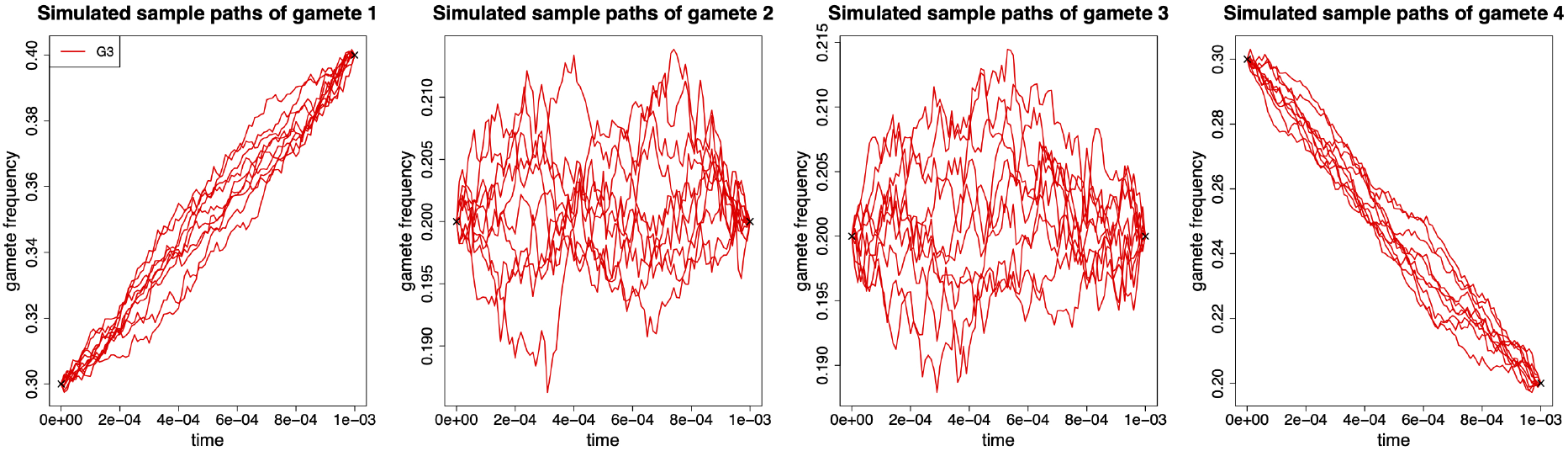
Simulated sample paths of each gamete evolving from generation 0 to 10 drawn from the *G*_*3*_-based importance sampler. In this illustration, we partition the time interval [0, 0.001] into 100 equidistant subintervals for the Euler-Maruyama scheme and generate 10 simulated sample paths with two observed states ***x***_0_ = (0.3, 0.2, 0.2, 0.3) and ***x***_0.001_ = (0.4, 0.2, 0.2, 0.2). Other parameters *N* = 5000, 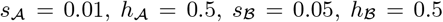 and *r* = 0.05.

Furthermore, the modification made in the *G*_*2*_-based importance sampler can be naturally applied to the *G*_*3*_-based importance sampler, which suggests setting ***ξ***_0_ = ***x***_0_ and ***ξ***_*U*_ = ***x***_*T*_, and successively drawing

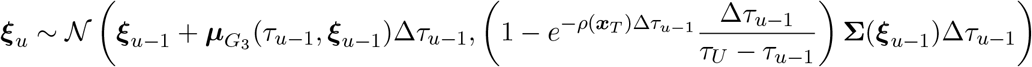

for *u* = 1, 2, … , *U* − 1. In Figure 9, we illustrate a set of simulated sample paths generated from the modified *G*_*3*_-based importance sampler with two observed states ***x***_0_ and ***x***_*T*_ as an example.

**Figure 9:**
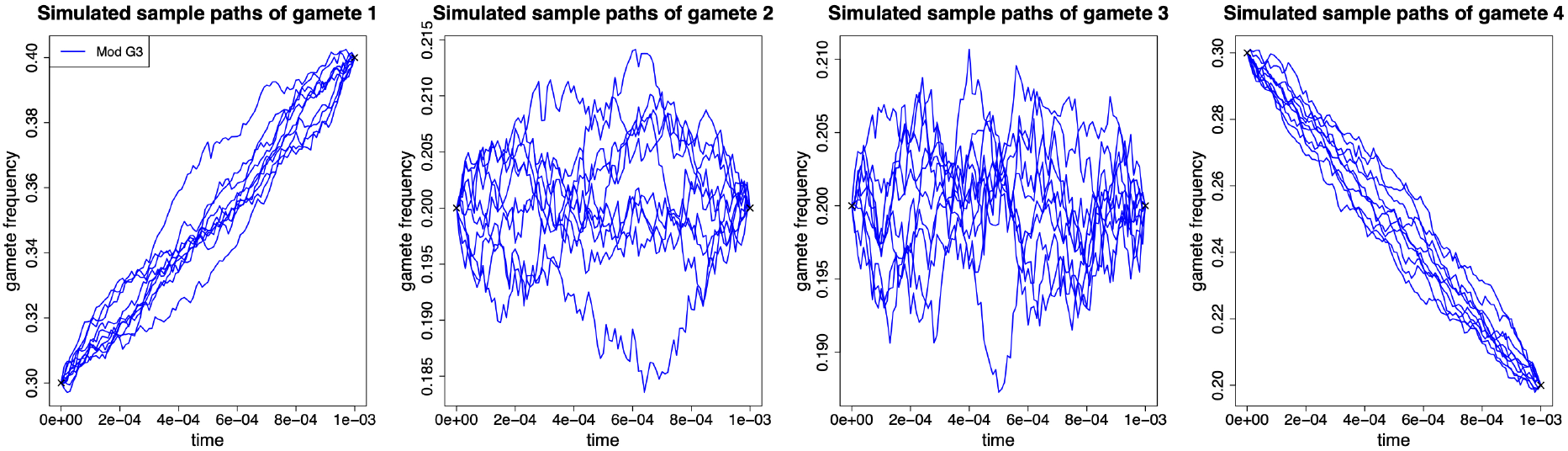
Simulated sample paths of each gamete evolving from generation 0 to 10 drawn from the modified *G*_*3*_-based importance sampler. In this illustration, we partition the time interval [0, 0.001] into 100 equidistant subintervals for the Euler-Maruyama scheme and generate 10 simulated sample paths with two observed states ***x***_0_ = (0.3, 0.2, 0.2, 0.3) and ***x***_0.001_ = (0.4, 0.2, 0.2, 0.2). Other parameters *N* = 5000, 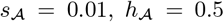, 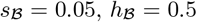 and *r* = 0.05.

Compared to the simulated sample paths generated from the Wright-Fisher diffusion shown in Figure 3, the simulated sample paths generated from the *G*-based importance samplers shown in Figures 5-9 more closely resemble typical paths that start at state ***x***_0_ and end at state ***x***_*T*_ since it is unlikely for the gamete frequency to undergo a drastic change at a specific time. This is due to the guiding drift term ***μ***_*G*_(*t*, ***x***) driving the simulated sample paths to terminate at the observed state ***x***_*T*_, which enables most of the realisations 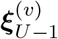 drawn from the *G*-based importance samplers to be in regions where the integrand 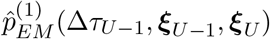 is large. The drift term ***μ***(***x***) of the Wright-Fisher diffusion incorporated into the *G* enables the simulated sample paths generated from the *G*-based importance samplers to imitate the local behaviour of the Wright-Fisher diffusion, which provides a sufficiently good approximation of the target. Hence, intuitively, the construction of the *G*-based importance samplers may result in much better performance.

Let 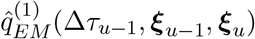 denote the probability density function of the multivariate normal distribution given in the *G*-based importance sampler and *w*(***ξ***_1_, ***ξ***_2_, … , ***ξ***_*U*−1_) denote the importance sampling weight given by

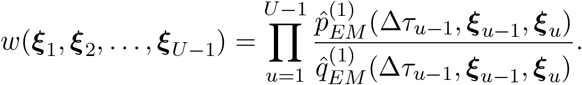

We can thereby approximate the Euler-Maruyama approximation 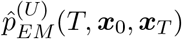 through importance sampling as

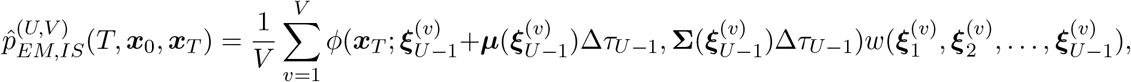

where 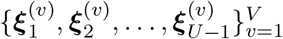 is the sample of the realisations successively generated from the *G*-based importance sampler, and *V* is the total number of the realisations in the sample. Similarly, we have

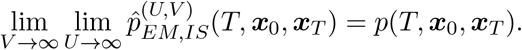

We illustrate the convergence of the importance sampling approximation 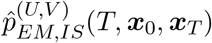 to the transition probability density *p*(*T*, ***x***_0_, ***x***_*T*_) on a logarithmic scale in Figure 10. Compared to the Monte Carlo approximation 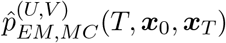 demonstrated in Figure 4, the variance resulting from Monte Carlo integration is largely removed through importance sampling with the *G*_*3*_-based importance sampler, especially for the modified *G*_*3*_ importance sampler.

**Figure 10:**
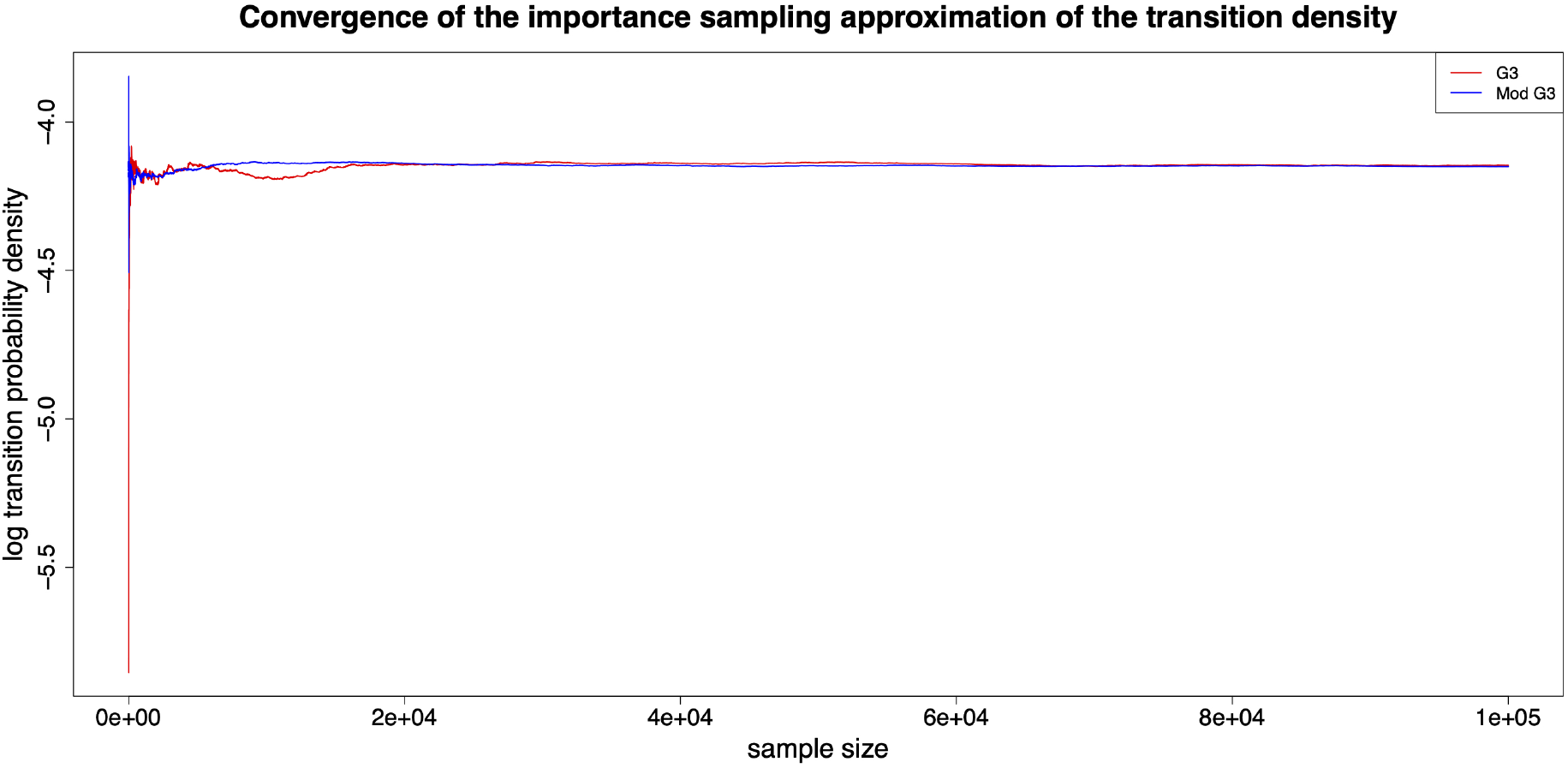
Convergence of the importance sampling approximation of the transition probability density 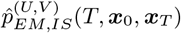 with the *G*_*3*_-based importance sampler. In this illustration, we partition the time interval [0, 0.001] into 100 equidistant subintervals for the Euler-Maruyama scheme and take ***x***_0_ = (0.3, 0.2, 0.2, 0.3), ***x***_0.001_ = (0.4, 0.2, 0.2, 0.2), *N* = 5000, 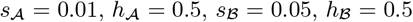 and *r* = 0.05.

**Figure 11:**
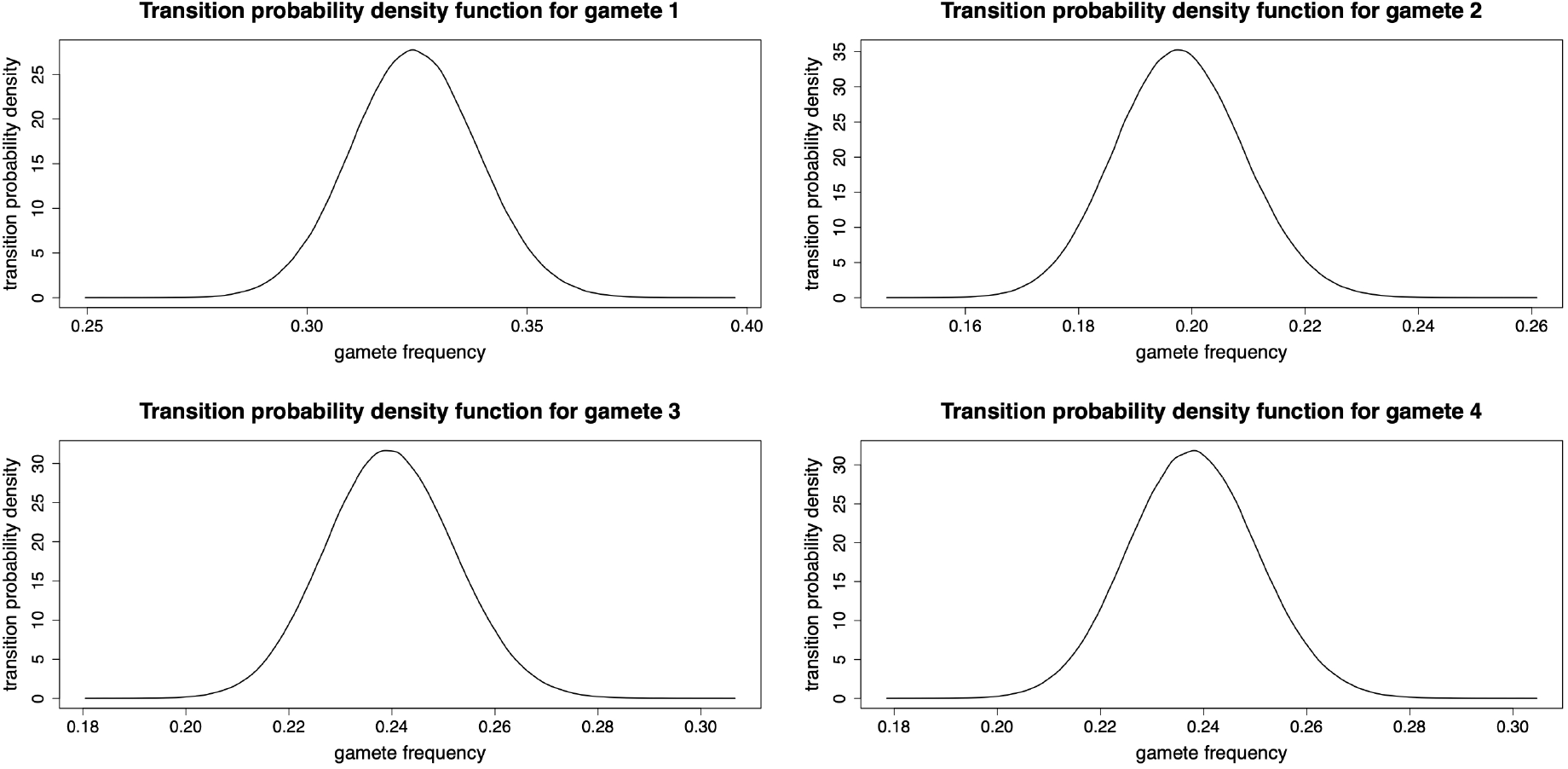
Transition probability density function for each gamete computed by kernel density estimation provided by a standard routine in the ks package of R. In this illustration, we partition the time interval [0, 0.001] into 100 equidistant subintervals for the Euler-Maruyama scheme and generate 10^6^ simulated sample paths starting from the state ***x***_0_ = (0.3, 0.2, 0.2, 0.3). Other parameters *N* = 5000, 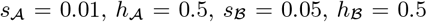 and *r* = 0.05.

### 4.4. Performance of the G-based Importance Samplers

We now evaluate the performance of the *G*-based importance samplers in the importance sampling approximation of the transition probability density 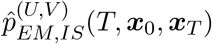, in particular for the *G*_*3*_-based importance sampler. Given that the explicit analytical transition density function of the Wright-Fisher diffusion is still unknown, we illustrate the convergence of the importance sampling approximation 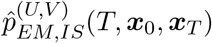 to the transition probability density *p*(*T*, ***x***_0_, ***x***_*T*_) with different *G*-importance samplers and evaluate their performance by looking at the effective sample size, denoted by *V*_*E*_, which means that the importance sampling approximation 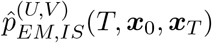 based on the weighted sample of size *V* is approximately as accurate as one based on an independent sample of size *V*_*E*_. From Liu (1996), the effective sample size is defined as

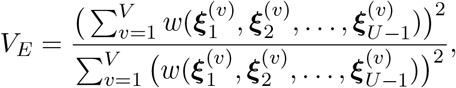

where 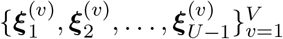 is the sample of the realisations generated from the *G*-based importance sampler, and *V* is the total number of the realisations in the sample.

We fix the observed initial state ***x***_0_ = (0.3, 0.2, 0.2, 0.3) and consider the observed terminal state ***x***_*T*_ for two cases: one is in the region where the underlying Wright-Fisher diffusion terminates at the time point *T* with a high probability, and the other is in the region where the underlying Wright-Fisher diffusion terminates at the time point *T* with a low probability. We take *T* = 0.001 and illustrate the transition probability density function for each gamete computed by kernel density estimation in Figure 3, which enable us to find high probability regions and low probability density regions where the underlying Wright-Fisher diffusion terminates at the time point *T* . From Figure 3, we pick ***x***_*T*_ = (0.32, 0.20, 0.24, 0.24) as the observed terminal state in a high probability region and ***x***_*T*_ = (0.38, 0.20, 0.24, 0.18) as the observed terminal state in a low probability region. We take use of a sample of the simulated sample paths generated from the proposal SDE of size *V* = 1000 to evaluate the importance sampling approximation 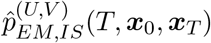 with different *G*-based importance samplers for these two observed terminal states, and show the convergence of the the importance sampling approximation 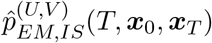 to the transition probability *p*(*T*, ***x***_0_, ***x***_*T*_) on a logarithmic scale in Figures 12 and 13, respectively. As shown in Figures 12 and 13, the *G*_*1*_-based importance sampler has the worst performance of all *G*-based importance samplers, and the *G*_*2*_- and *G*_*3*_-based importance samplers have quite similar performance.

**Figure 12:**
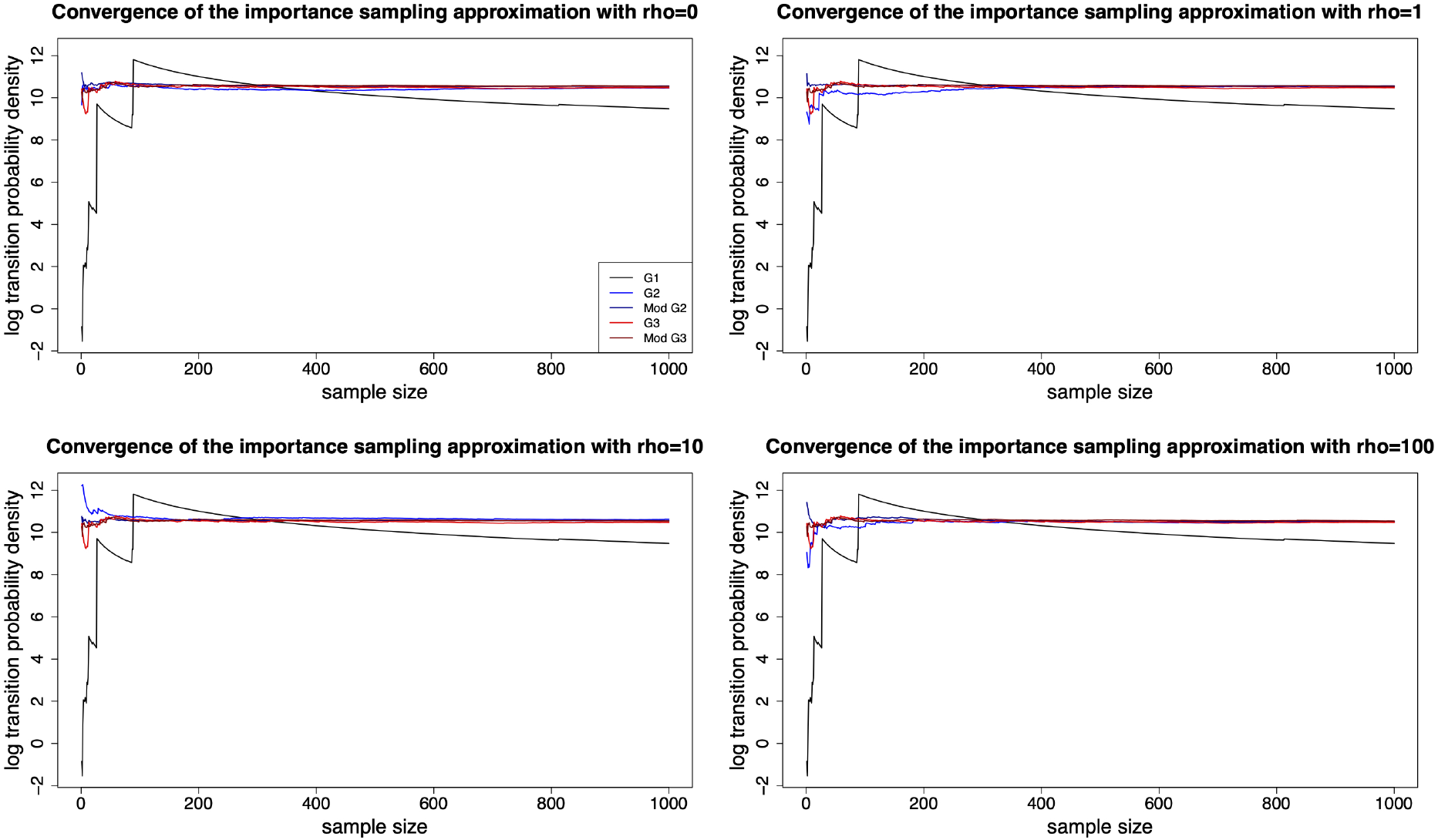
Comparison of the convergence of the importance sampling approximation of the transition probability density 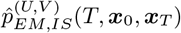 with different *G*-based importance samplers for the observed terminal state in a high probability region. In this illustration, we partition the time interval [0, 0.001] into 100 equidistant subintervals in the Euler-Maruyama scheme and generate 10^3^ simulated sample paths with two observed states ***x***_0_ = (0.3, 0.2, 0.2, 0.3) and ***x***_0.001_ = (0.32, 0.20, 0.24, 0.24). Other parameters *N* = 5000, 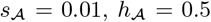, 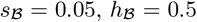 and *r* = 0.05.

**Figure 13:**
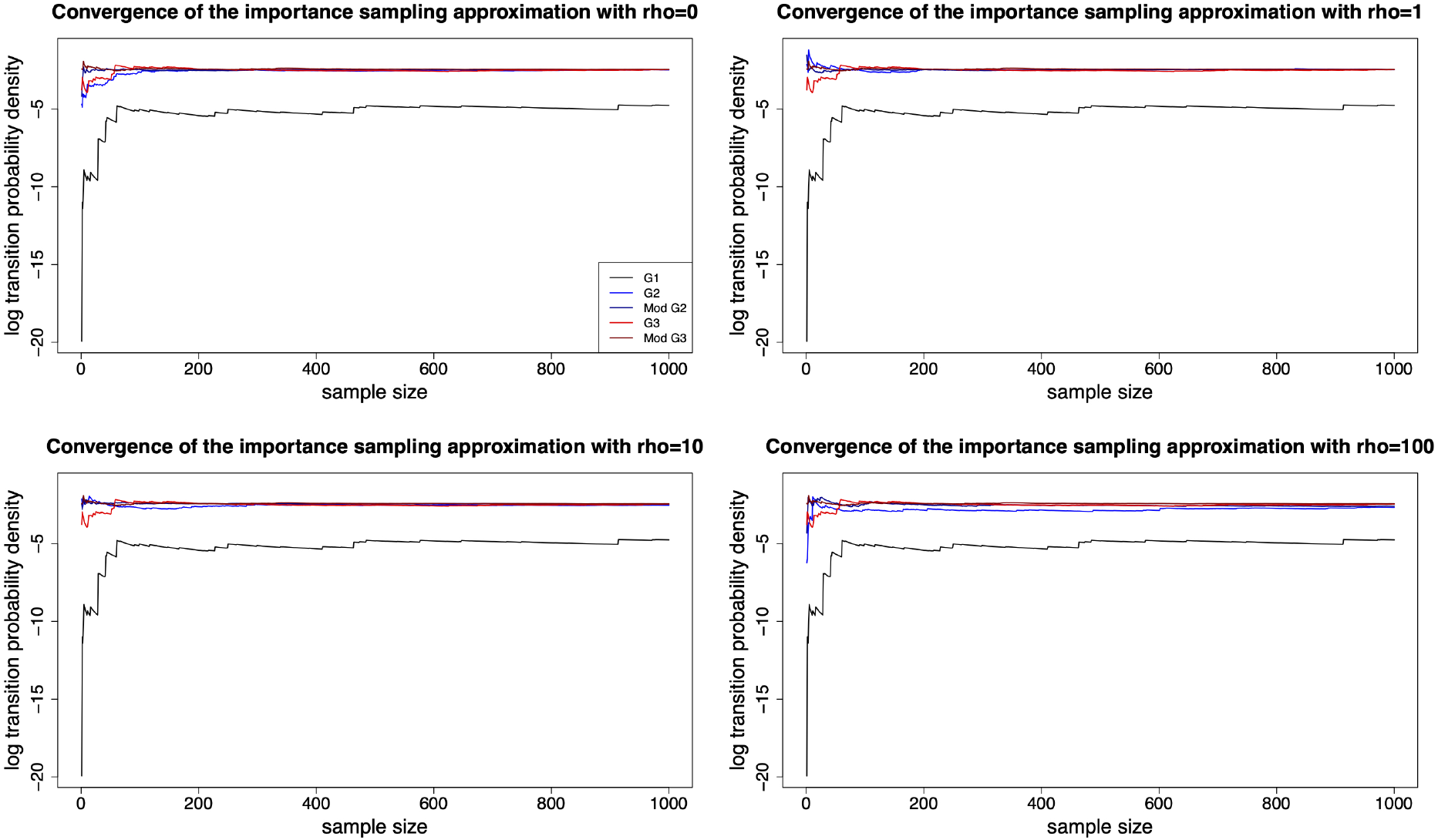
Comparison of the convergence of the importance sampling approximation of the transition probability density 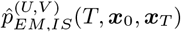 with different *G*-based importance samplers for the observed terminal state in a low probability region. In this illustration, we partition the time interval [0, 0.001] into 100 equidistant subintervals in the Euler-Maruyama scheme and generate 10^3^ simulated sample paths with two observed states ***x***_0_ = (0.3, 0.2, 0.2, 0.3) and ***x***_0.001_ = (0.38, 0.20, 0.24, 0.18). Other parameters *N* = 5000, 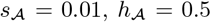, 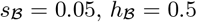 and *r* = 0.05.

For further comparison of the performance of the *G*-based importance samplers, we replicate the importance sampling procedure 100 times and evaluate the average effective sample size for importance sampling based on different *G*-based importance samplers in Table 1. As shown in Table 1, the *G*_*1*_-based importance sampler has the smallest effective sample size for importance sampling of all *G*-based importance samplers, which confirms the results in Figures 12 and 13. Overall, the effective sample size is larger for importance sampling based on the *G*_*3*_-based importance sampler than other *G*_*2*_-based importance samplers, especially for the modified *G*_*3*_-based importance sampler. For the observed state ***x***_*T*_ in a high probability region, the effective sample size is larger for importance sampling based on the *G*_*2*_-based importance sampler with small values of the *ρ*-term than large values. In contrast, the effective sample size is larger for importance sampling based on the *G*_*2*_-based importance sampler with large values of the *ρ*-term than small values for the observed state ***x***_*T*_ in a low probability region. These results confirm our intuition in the design of the *ρ*-term to improve the performance of the *G*_*3*_-based importance sampler, *i.e.*, it adjusts the contribution of the drift term of the underlying Wright-Fisher diffusion and the guiding drift term on the proposal SDE according to the observed terminal state ***x***_*T*_ . Moreover, from Table 1, we find that the effective sample size is larger for importance sampling based on the modified *G*_*3*_-based importance sampler than the original one, which implies that the modification suggested by Fearnhead (2008) works for the *G*_*3*_-based importance sampler. In summary, our results indicate that the modified *G*_*3*_-based importance sampler has the best performance of all *G*-based importance samplers.

**Table 1:** Average effective sample size for the importance sampling approximation of the transition probability density 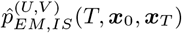 based on different *G*-based importance samplers.

| Importance sampler | Average effective sample size |  |
| --- | --- | --- |
| | $\mathbf{x}_T = (0.32, 0.20, 0.24, 0.24)$ | $\mathbf{x}_T = (0.38, 0.20, 0.24, 0.18)$ |
| $G_1$ | 4.727507 | 5.510372 |
| $G_2$ with $\rho = 0$ | 17.262143 | 13.079748 |
| $G_2$ with $\rho = 1$ | 19.262232 | 12.439503 |
| $G_2$ with $\rho = 10$ | 20.237910 | 11.992605 |
| $G_2$ with $\rho = 100$ | 19.078593 | 10.780667 |
| $G_2$ (modified) with $\rho = 0$ | 18.211303 | 14.126935 |
| $G_2$ (modified) with $\rho = 1$ | 20.677225 | 12.980928 |
| $G_2$ (modified) with $\rho = 10$ | 21.696237 | 12.597953 |
| $G_2$ (modified) with $\rho = 100$ | 19.456650 | 11.723928 |
| $G_3$ | 19.440782 | 13.301470 |
| $G_3$ (modified) | 20.459668 | 15.731508 |

To further improve the importance sampling approximation, we can adopt the asymptotically optimal choice of the total number of the realisations in the sample 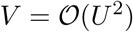 to balance the trade-off between reducing the bias caused by the Euler-Maruyama scheme and reducing the variance resulting from Monte Carlo integration (Stramer & Yan, 2007), and take the use of antithetic variates for variance reduction, which is applied in Brandt & Santa-Clara (2002). There are other acceleration techniques, bias reduction techniques and variance reduction techniques discussed in Durham & Gallant (2002).

## 5. Discussion

In this work, we constructed two structurally different Wright-Fisher diffusion driven SDEs to characterise the population dynamics under natural selection at two linked loci in terms of the changes in gamete frequencies over time. One of these two SDEs has closed-form expressions for both of the drift term and the diffusion term, which can contribute to an efficient numerical sample path solution for the SDE satisfied by the two-locus Wright-Fisher diffusion with selection. Alternatively, we can describe the population evolving under natural selection at two linked loci in terms of the changes in allele frequencies and linkage disequilibrium over time with the Wright-Fisher diffusion driven SDE constructed in Appendix B, which however is of a more complicated form. We therefore only provided the derivation of the Wright-Fisher diffusion in Appendix B.

We developed an efficient importance sampling approximation of the transition probability density of the Wright-Fisher diffusion between two observed states. We evaluated the performance of several guided process based importance samplers. The main idea behind the design of the guiding term is to enable the sample path of the proposal SDE to hit the observed states while at the same time imitate the local behaviour of the underlying Wright-Fisher diffusion. Our importance sampler is an extension of the proposal introduced by Fearnhead (2008), where we take the *ρ*-term in the proposal of Fearnhead (2008) to be a function of the observed state ***x***_*T*_ . Such a modification is used to adjust the contributions of the drift term of the underlying Wright-Fisher diffusion and the guiding drift term on the proposal SDE according to the observed state ***x***_*T*_, so that the resulting proposal SDE can evolve largely under the influence of the underlying Wright-Fisher diffusion if its sample path is already close to the given state ***x***_*T*_ with a high probability, but allow a more significant contribution from the guiding term if its sample path gets further away from the given state ***x***_*T*_ with a high probability. A simple comparison suggests that our importance sampler may have better performance, which can be further improved through various acceleration techniques, bias reduction techniques and variance reduction techniques proposed by Durham & Gallant (2002).

In recent years, the Wright-Fisher diffusion has already been successfully applied in the population genetic analysis of time series data of allele frequencies, but most of these methods are limited to either a single locus or multiple independent loci, *i.e.*, genetic recombination effect and local linkage information are ignored in these methods. However, in some cases accounting genetic recombination and linkage information is essential for the inference of natural selection, especially when two loci are tightly linked. Otherwise incorrect results might be obtained with existing single-locus methods (see Figure 14 and 15). Our method can play an important role in the development of the many existing simulation-based inference approaches such as simulated maximum-likelihood methods and Markov chain Monte Carlo methods for the inference of natural selection from generic time series data

**Figure 14:**
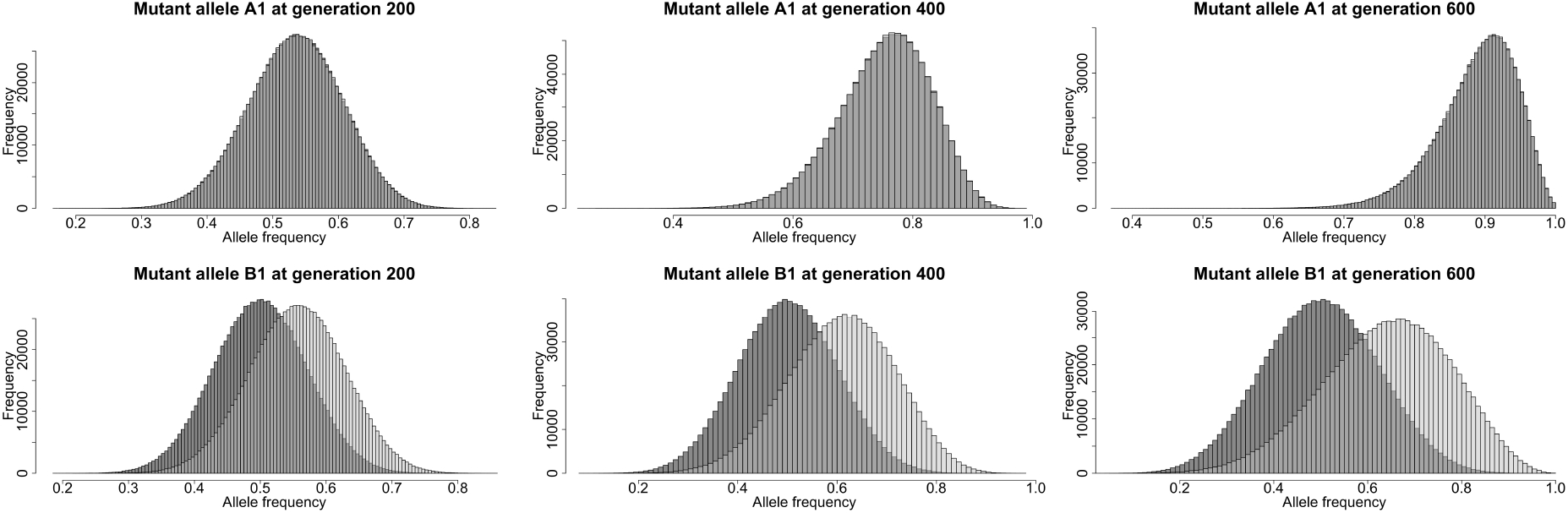
A example of a positively selected locus 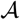 tightly linked with a neutral locus 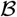. The initial haplotype frequencies of the population are ***x***_0_ = (0.2, 0.1, 0.3, 0.4), and the selection coefficients are 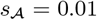 and 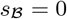. Dark grey denotes the one-locus Wright-Fisher diffusion, and light grey denotes the the two-locus Wright-Fisher diffusion.

**Figure 15:**
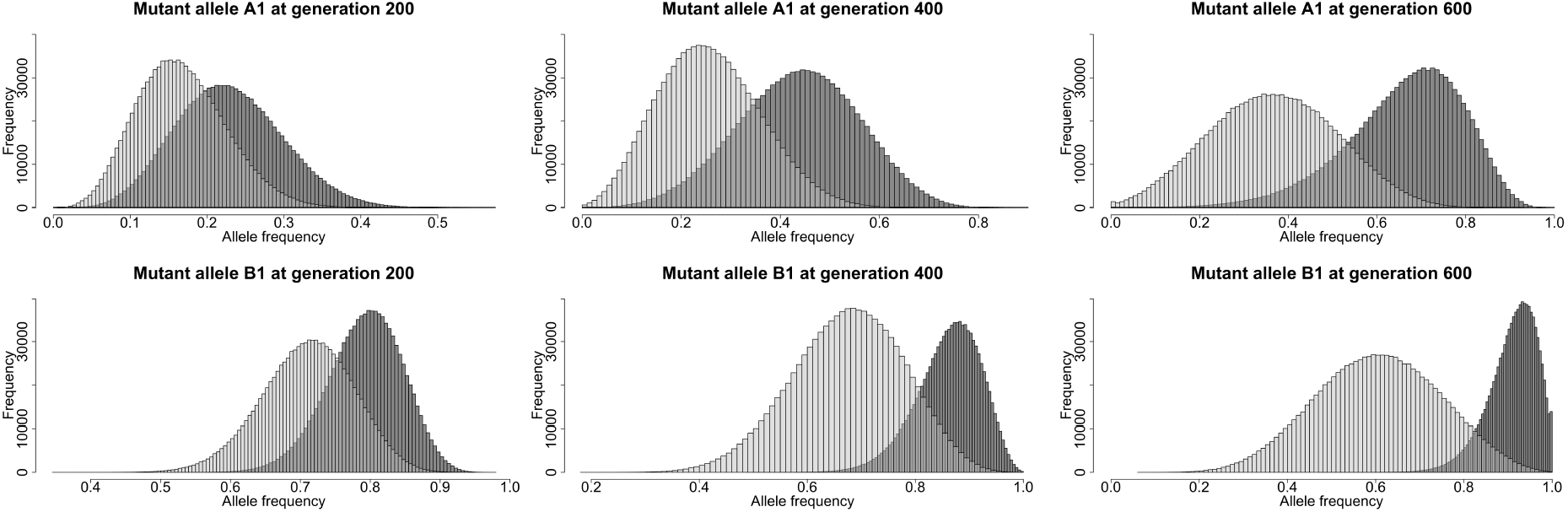
A example of two tightly linked loci 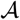 and 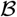 under natural selection. The initial haplotype frequencies of the population are ***x***_0_ = (0.05, 0.05, 0.7, 0.2), and the selection coefficients are 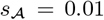 and 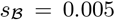. Dark grey denotes the one-locus Wright-Fisher diffusion, and light grey denotes the the two-locus Wright-Fisher diffusion.

Given that many phenotypes are known to show variation caused by multiple linked loci, there is strong motivation to consider natural selection at more than one locus. However, theoretical studies of natural selection simultaneously acting on multiple linked loci are much more challenging. Even for the case of two linked loci, the population evolving subject to natural selection is complicated and has not been completely understood. According to Hamilton (2009), genetic recombination promotes linkage equilibrium whereas natural selection maximises the mean fitness, and the equilibrium is determined by the relative strength of these two mechanisms. When the two processes are of approximately equal strength, the result is a compromise that may produce an equilibrium that is neither in linkage equilibrium nor at maximum mean fitness. In addition, the equilibrium also depends strongly on gene actions (*i.e.*, the dominance parameters 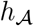 and 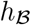) and the initial state of the population. However, all of these conclusions have been reached under the assumption of the infinite effective population size, which is quite restrictive and is never met in natural populations. In this section, we investigate the effect of natural selection acting on the population dynamics at two linked loci based on the simulated Wright-Fisher diffusion sample paths.

We illustrate several examples of the population dynamics subject to natural selection at two linked loci in Figures 16-20, where we partition the time interval into 5,000 equidistant subintervals in the Euler-Maruyama scheme and evaluate the empirical transition probabilities with 1,000,000 independent realisations of the Wright-Fisher diffusion. Figures 16 and 17 demonstrate the cases of the population initially in positive and negative linkage disequilibrium, respectively. Figure 18 gives an example when the positively selected allele is initially in low frequency. Figure 19 shows an example of non-additive gene action. Figure 20 demonstrates the case when only locus 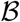 is subject to natural selection. As illustrated in Figures 16-20, we find that genetic recombination promotes linkage equilibrium whereas natural selection promotes the fixation of the gamete type favoured by natural selection, 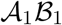, *i.e.*, the type with the highest mean fitness, which confirms the results stated in Hamilton (2009). However, in finite populations, linkage disequilibrium always decays to zero, *i.e.*, linkage equilibrium is eventually reached. When both of the two loci are positively selected, natural selection leads eventually to the fixation of a certain gamete type 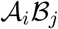, but not necessarily the 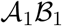 gamete. In fact, which gamete type becomes fixed is a random event with a corresponding fixation probability determined by the population size, the strength of natural selection and genetic recombination, gene actions and the initial state of the population (see Figures 16-19). By contrast, when at least one of the two loci is neutral, the fixation of a certain gamete type 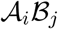 may not occur (see Figure 20).

**Figure 16:**
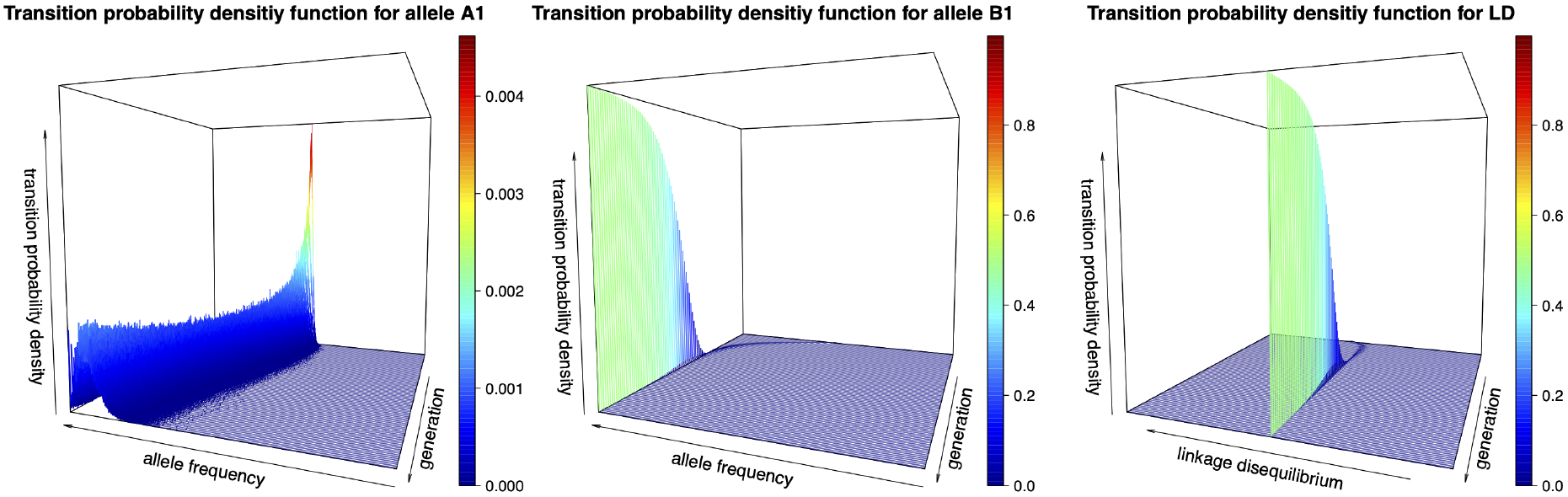
Dynamics of the empirical transition probabilities for the 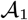 allele, the 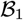 allele and linkage disequilibrium over 500 generations for the population initially in positive linkage disequilibrium. We take other parameters to be ***x***_0_ = (0.3, 0.2, 0.2, 0.3), *N* = 5000, 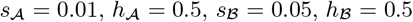 and *r* = 0.05.

**Figure 17:**
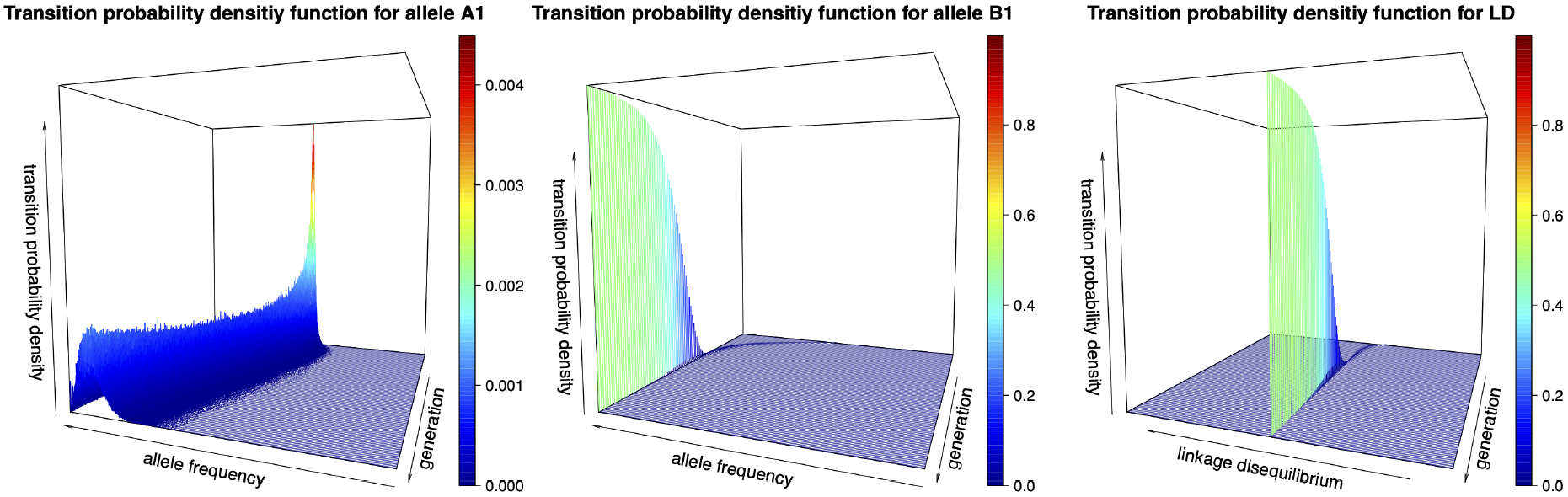
Dynamics of the empirical transition probabilities for the 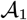 allele, the 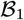 allele and linkage disequilibrium over 500 generations for the population initially in negative linkage disequilibrium. We take other parameters to be ***x***_0_ = (0.2, 0.3, 0.3, 0.2), *N* = 5000, 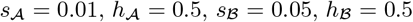 and *r* = 0.05.

**Figure 18:**
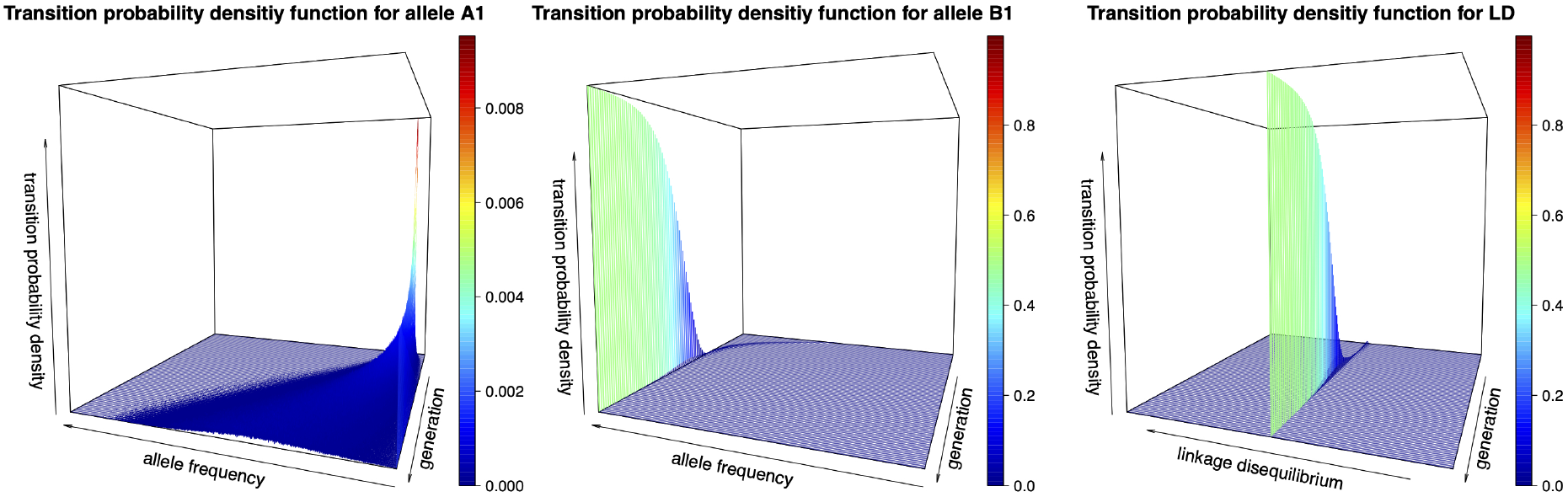
Dynamics of the empirical transition probabilities for the 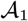 allele, the 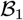 allele and linkage disequilibrium over 500 generations for the allele favoured by natural selection initially in low frequency. We take other parameters to be ***x***_0_ = (0.03, 0.02, 0.48, 0.47), *N* = 5000, 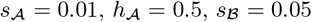 and *r* = 0.05.

**Figure 19:**
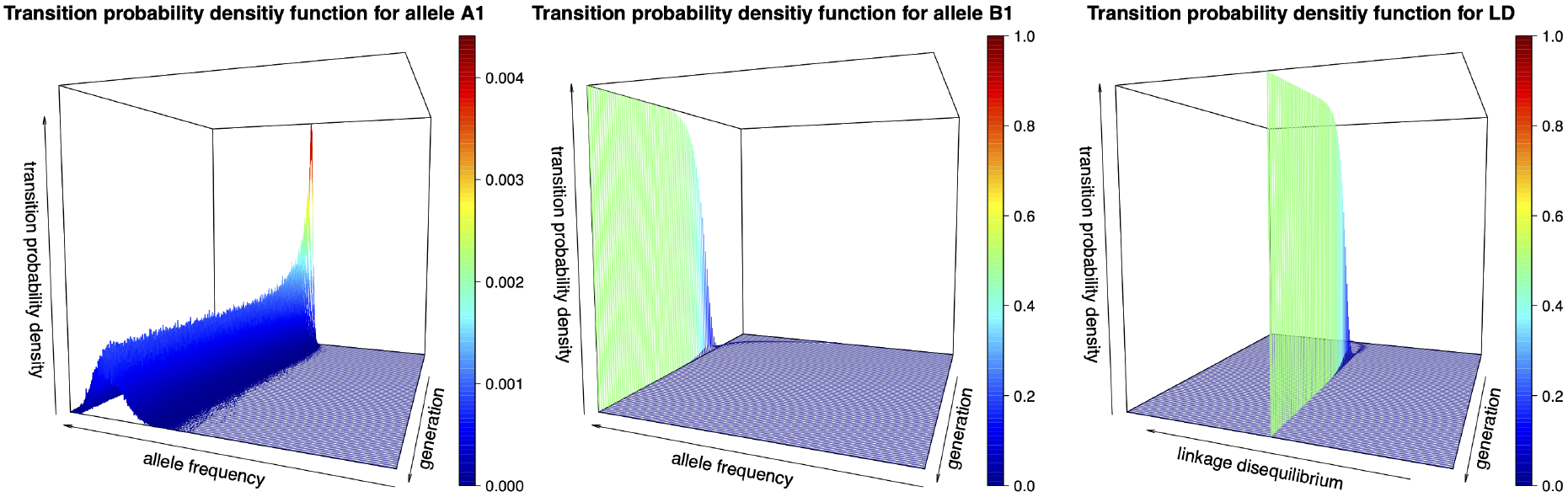
Dynamics of the empirical transition probabilities for the 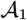 allele, the 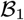 allele and linkage disequilibrium over 500 generations for non-additive gene actions. We take other parameters to be ***x***_0_ = (0.3, 0.2, 0.2, 0.3), *N* = 5000, 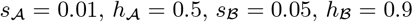 and *r* = 0.05.

**Figure 20:**
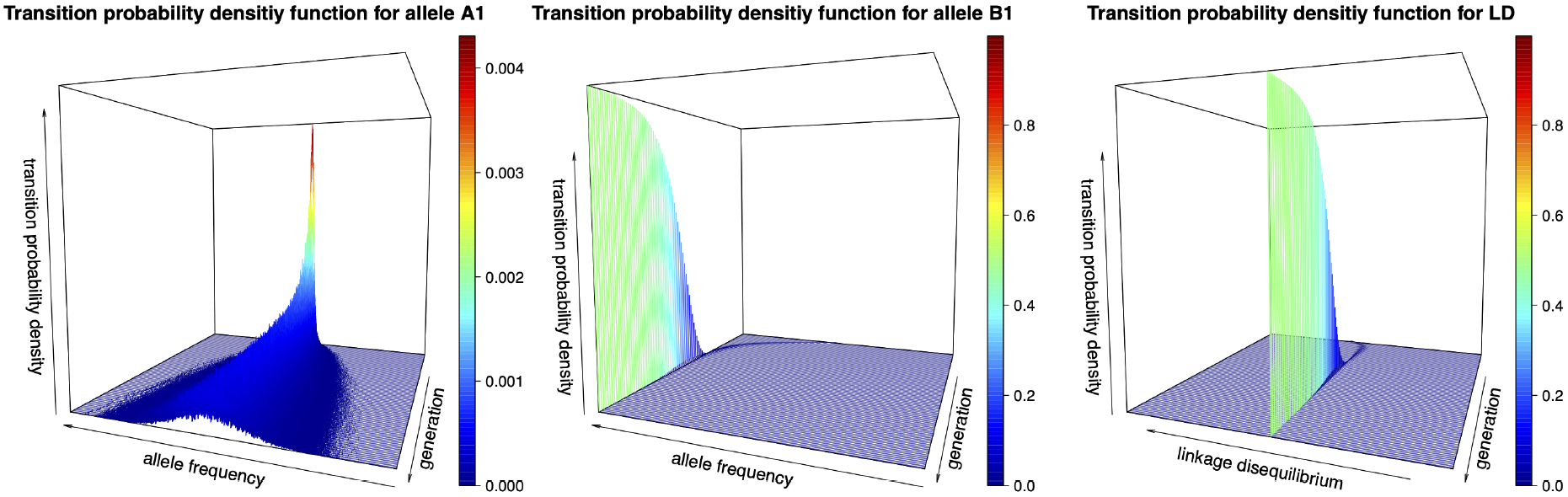
Dynamics of the empirical transition probabilities for the 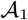 allele, the 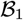 allele and linkage disequilibrium over 500 generations for neutral locus 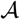. We partition the time interval into 5000 equidistant subintervals in the Euler-Maruyama scheme and evaluate the empirical transition probabilities with 10^6^ independent realisations of the Wright-Fisher diffusion. We take other parameters to be ***x***_0_ = (0.3, 0.2, 0.2, 0.3), *N* = 5000, 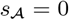, 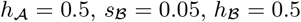 and *r* = 0.05.

The scope of our results in the present work can be extended in various directions. A direct extension is to certain multi-locus problems. As we have stated above, in theory, our method can be naturally extended to high-dimensional problems, but may suffer from the potential issue of computational efficiency in practice. We have two potential directions to improve the performance of our method for multi-locus problems. One direction is to consider how to exactly simulate the sample path of the Wright-Fisher diffusion to reduce the bias caused by numerical simulation schemes in this work. A series of recent papers (*e.g.*, Jenkins & Spanó, 2017) may provide some new inspirations. The other direction is to consider how to construct a more proper proposal SDE in importance sampling. In this work, we simply apply the Mahalanobis distance (De Maesschalck et al., 2000) to construct the *ρ*-term in our importance sampler, which roughly measures the distance from the observed state ***x***_*T*_ to the regions where the underlying Wright-Fisher diffusion sample path terminates at time *T* with a high probability. A better construction of the *ρ*-term may improve the performance of the importance sampling approximation of the transition probability density.

## Acknowledgements

This work was funded in part by the Engineering and Physical Sciences Research Council (EPSRC) Grant EP/I028498/1 to F.Y.

## Appendix A. Proof of Theorem 2

Let 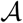 denote the generator associated with the SDE in Eq. (4), then

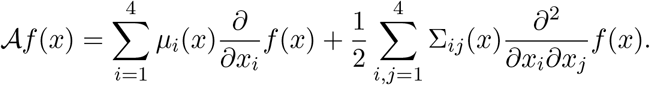

Let 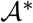 be the generator associated with the SDE in Eq. (12), then

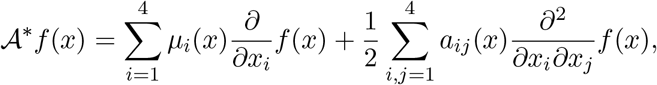

where

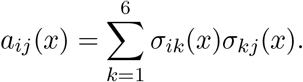

From (13), we have

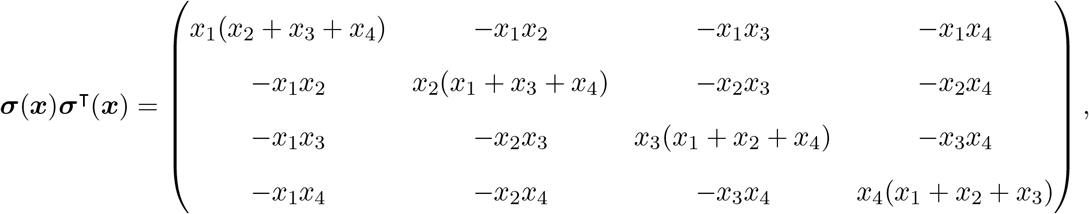

which is the same as Σ_*ij*_(***x***) = *x*_*i*_(*δ*_*ij*_ − *x*_*j*_) given in (7). This shows the two SDE’s in Eqs. (4) and (12) have the same weak solutions.

## Appendix B. An alternative formulation of the Wright-Fisher diffusion

We provide an alternative formulation of the Wright-Fisher diffusion for the population dynamics under natural selection at two linked loci. From Eq. (8), it is clear that the population evolving according to the Wright-Fisher diffusion ***X*** can be characterised by considering any three linearly independent gamete frequencies out of the four, *e.g.*, *X*_1_, *X*_2_ and *X*_3_. However, there is an obvious asymmetry in doing so, and based on Ewens (2004), we find it more convenient to work with a vector of new independent variables 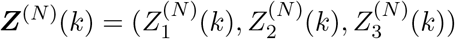, which is related to ***X***^(*N*)^(*k*) in the following way:

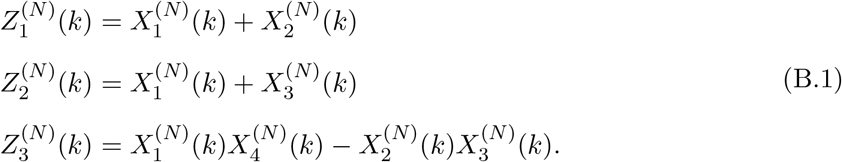

Note that 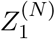 is the frequency of the 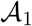 allele, 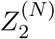 is the frequency of the 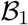 allele, and 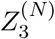 is the standard measure of linkage disequilibrium.

### Theorem 3.

*If we let the population size N go to infinity and hold the scaled parameters* 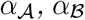 *and R constant, the Markov process of mutant allele frequencies and linkage disequilibrium **Z***^(*N*)^ *converges to a time-homogeneous diffusion process, denoted by **Z*** = *{**Z***(*t*), *t* ≥ 0}, *satisfying the following stochastic differential equations of the form*

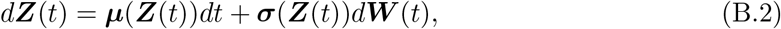

*where the drift term **μ***(***z***) *is a three-dimensional vector being*

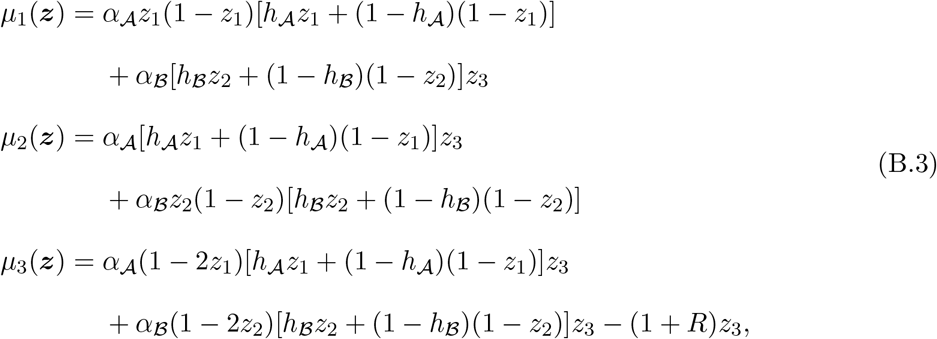

*the diffusion term **σ***(***z***) *is three by three matrix being the square root of the infinitesimal covariance matrix* **Σ**(***z***) *given by*

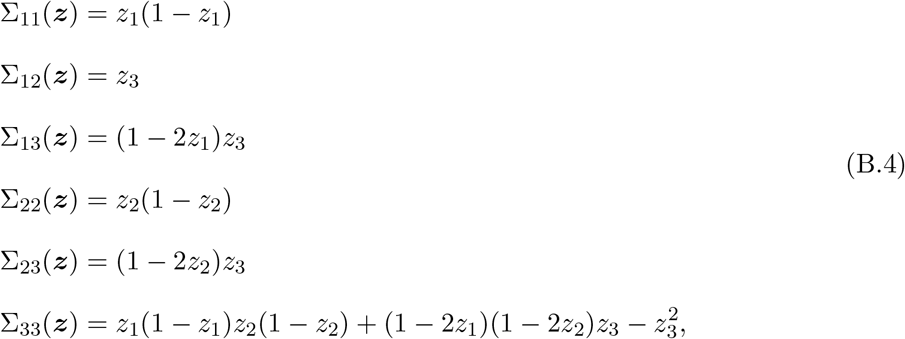

*and **W*** (*t*) *is a three-dimensional standard Brownian motion.*

The proof of Theorem 3 is almost identical to the proof for neutral populations in Durrett (2008, p. 324) by using a multivariate extension of the change of variables formula. We refer to the process of mutant allele frequencies and linkage disequilibrium ***Z*** solving the SDE in Eq. (B.2) as the two-locus Wright-Fisher diffusion with selection, which evolves in the state space

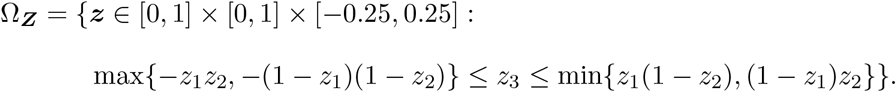

From Eqs. (B.3) and (B.4), we find that the drift term ***μ***(***z***) and the diffusion term ***σ***(***z***) satisfy both global Lipschitz and linear growth conditions (8) and (9) for all ***z*** ∈ Ω_***Z***_, which guarantees the existence and uniqueness of a non-explosive solution of the Wright-Fisher diffusion driven SDE for any given ***Z***(0) independent of ***W*** (*t*) and satisfying *E*(|***Z***(0)|^2^) < ∞.

Moreover, for any ***z*** in the interior of the state space Ω_***Z***_, the infinitesimal mean vector ***μ***(***z***) and the infinitesimal covariance matrix **Σ**(***z***) are twice continuously differentiable with respect to ***z*** with bounded derivatives, and the infinitesimal covariance matrix **Σ**(***z***) is also positive definite, which implies that there exists a well-defined and smooth transition probability density function of the Wright-Fisher diffusion ***Z*** satisfying the Kolmogorov backward equation

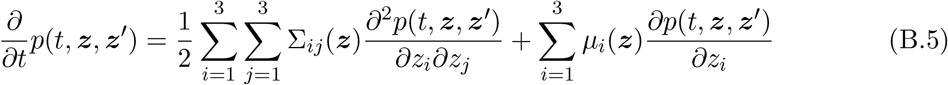

for fixed ***z′*** and the Kolmogorov forward equation

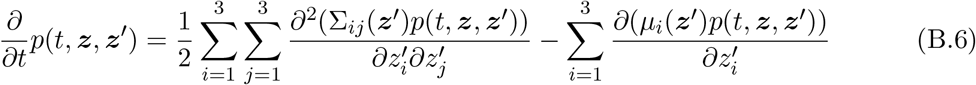

for fixed ***z***, where the infinitesimal mean vector ***μ***(***z***) and the infinitesimal covariance matrix **Σ**(***z***) are separately given by Eqs. (B.3) and (B.4).

## Notes

### Competing Interest Statement

The authors have declared no competing interest.

